# Multiomics Reveals Compartmentalized Immune Responses and Tissue-Vascular Signatures in Lyme Disease

**DOI:** 10.1101/2025.05.27.656061

**Authors:** Clifford Rostomily, Ameek Bhalla, Henry Hampton, Lance Pflieger, Mary E. Brunkow, Denise Cooper, Christopher Lausted, Ajay Suresh Akhade, Brett Smith, Kelsey Scherler, Pamela Troisch, Kai Wang, Gary P. Wormser, Noa Rappaport, Leroy Hood, Naeha Subramanian

**Affiliations:** Institute for Systems Biology, Seattle, WA, USA; Division of Infectious Diseases, New York Medical College, Valhalla, New York, USA; Department of Immunology, University of Washington, Seattle WA, USA; Phenome Health, Seattle, WA, USA; Buck Institute for Research on Aging, Novato, CA, USA

**Keywords:** Lyme disease, *Borrelia burgdorferi*, PBMC, skin, endothelia, multiomics, immune response

## Abstract

Lyme disease (LD) is growing in incidence, with nearly 500,000 cases diagnosed annually in the United States. Despite treatment, some patients experience persistent symptoms. The immune mechanisms underlying LD remain poorly understood. We conducted a multiomic longitudinal analysis of 49 LD patients and matched controls, integrating plasma proteomics, metabolomics, PBMC immunophenotyping, and a meta-analysis of skin lesions. We identified compartmentalized immune responses in acute LD, with coordinated alterations in circulating plasma proteins and metabolites linked to endothelial barrier stability, metabolic reprogramming, and symptom severity, predominantly traced to tissue and vascular immune processes at the site of infection. In contrast, PBMCs remained largely quiescent, revealing a disconnect between localized tissue responses and systemic immunity. These findings provide novel insights into LD pathophysiology and highlight the potential for diagnostics leveraging tissue and vascular immune markers detectable in blood. They also provide a resource for biomarker discovery and predictive modeling to improve LD management.

## Introduction

Lyme Disease (LD) is the most prevalent vector-borne disease in Europe and North America, with an estimated 476,000 new cases diagnosed annually in the United States alone^1^. The principal causative agent in the United States, *Borrelia burgdorferi* sensu stricto (*B. burgdorferi* s.s.), is a bacterial spirochete transmitted to humans most often through the bite of infected *Ixodes scapularis* ticks. Primarily residing in its main reservoir host, the white-footed mouse (*Peromyscus leucopus*), the spirochete employs sophisticated immune evasion strategies^2^ and persists in the reservoir host with minimal impact on host survival^3^. Transmission to humans occurs by a tick bite most often during the spring and summer months, when ticks are most active. The geographic expansion of LD, impacted by climate change and the proliferation of suitable tick habitats, has led to its emergence in previously unaffected areas^4,5^.

LD clinical manifestations are categorized as early localized, early disseminated, and late disseminated stages. During the early localized stage, 70–95% of infected patients develop erythema migrans (EM), an erythematous circular rash that develops at the site of the tick bite, typically 7–10 days post-infection^6,7^. This stage is often accompanied by flu-like symptoms such as fever, fatigue, headache, and joint pain^8^. In the early disseminated stage, the pathogen spreads hematogenously to other sites, most often causing additional EM skin lesions, but also potentially carditis, and certain neurologic manifestations such as facial nerve palsy. If untreated, LD may progress to the late disseminated stage, characterized most often in the United States by large joint arthritis^9^. While antibiotic treatment is effective in most cases, 10–20% of patients experience persistent subjective symptoms lasting six months or longer, termed Post Treatment Lyme Disease Syndrome (PTLDS) if they are severe enough to be disabling^10,11^. PTLDS is defined by a heterogeneous mix of symptoms, including fatigue, cognitive difficulties, and musculoskeletal pain, with an unclear etiology. In certain aspects PTLDS is similar to long COVID^12^. In view of the fact that LD may affect multiple different organ systems, as reflected in its diverse clinical manifestations, understanding the host immune response is crucial for identifying biomarkers that could improve diagnostics, facilitate early intervention, and mitigate long-term impacts.

Despite its prevalence, LD lacks an approved vaccine for humans, and current diagnostic methods have significant limitations. Standard serologic tests, such as an ELISA and Western blot, which detect host antibodies to *B. burgdorferi*, often remain negative for several weeks after the onset of infection, limiting their clinical utility. Blood culture and PCR-based tests on blood are limited by relatively low sensitivity due to the low abundance and/or frequency of *B. burgdorferi* in blood samples^13^. Two-tier serologic testing has low sensitivity in patients with EM (the most common clinical manifestation of LD) and cannot distinguish active infection from past exposure. Variability in humoral immune responses among individuals further undermines the reliability of antibody-based diagnostics^14-16^. Additionally, *B. burgdorferi* actively disrupts host immune responses through strategies such as antigenic variation and impairing B cell maturation, germinal center formation, and the production of protective antibodies and memory cells^17-19^. While localized interferon responses and systemic Th1 cytokines (e.g., IFN-γ) have been observed in human LD^20,21^, these responses appear insufficient for pathogen clearance and may even facilitate dissemination in murine models^22^. These challenges underscore the need for an integrated understanding of the host response across tissues, immune compartments, and metabolic systems to inform novel diagnostic and therapeutic strategies.

In this study, we performed a multiomic longitudinal analysis of 49 LD patients and matched controls. By integrating data from 460 plasma proteins, 1244 metabolites, deep immunophenotyping of PBMCs, and a meta-analysis of skin lesions, we provide a comprehensive systems-level view of the immune response to LD. Our findings reveal that tissue and vascular immune processes dominate the host response, driving significant alterations in circulating proteins and metabolites, while peripheral blood immune cells remain relatively quiescent. This critical disconnect between localized tissue responses and systemic immunity offers new insights into the pathophysiology of LD, with implications for diagnostics based on tissue and vascular immune markers detectable in the blood. As the first multiomic analysis of LD at this scale, this work provides a valuable resource for the scientific community and lays the groundwork for future research aimed at predictive modeling of LD progression and improved clinical management of the disease.

## Results

### Study overview and clinical manifestations

A longitudinal cohort of 49 adult LD patients and age- and sex-matched healthy controls was recruited at the Lyme Disease Diagnostic Center (LDDC), New York Medical College. Peripheral blood mononuclear cells (PBMCs) and plasma were collected from patients at four time points: diagnosis (T1), 2–3 weeks post-diagnosis (T2), 6 months (T3), and 1 year (T4). Control samples were collected at T1, T3, and T4 to account for possible seasonal variations in immune parameters **(Fig. 1a)**. The demographic composition of the cohort is shown in **Fig. 1b**. All patients were treated with antibiotics following their initial blood draw, primarily doxycycline, although some patients received alternative regimens based on physician recommendations **(Table 1)**. At recruitment, a subset of patients (n=28) had already started antibiotic treatment, ranging from 0–28 days, after visiting a primary care physician before referral to the LDDC. These patients were categorized as antibiotic non-naïve, while the remaining 22 patients who had not yet received antibiotics were classified as antibiotic-naïve **(Fig. S1a)**.

**Figure 1:**
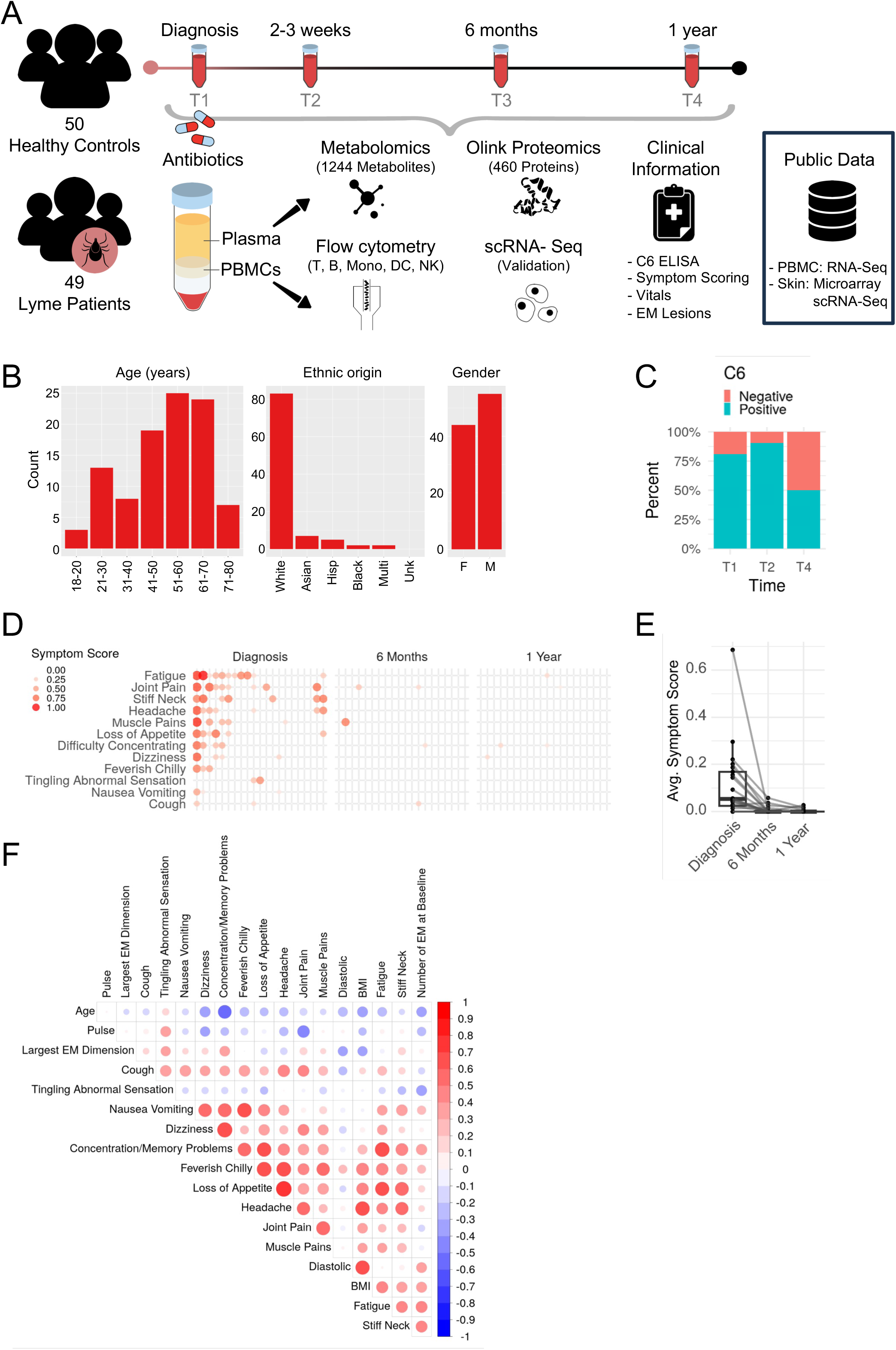
Study overview and clinical manifestations of Lyme disease (LD). **(A)** Schematic of the study. Blood was drawn at four time points from age- and sex-matched patients (n=49) and controls (n=50). Blood samples were fractionated into plasma and PBMCs on which the indicated assays were performed. Detailed clinical information and symptom questionnaires were collected from the patients. Public transcriptomic data of PBMCs and erythema migrans (EM) lesions from LD patients were also analyzed in parallel. **(B)** Demographic characteristics of the cohort. Bar plots show the total number of individuals in the study cohort, categorized by age, sex (F: female; M: male), and ethnicity (Hisp: Hispanic; Multi: Multiethnic, Unk: Unknown). **(C)** Stacked bar chart showing the proportion of antibiotic-naive patients that tested positive for LD by C6 ELISA at diagnosis (T1), 6 months (T2), and 1 year (T4). **(D)** Bubble plot showing patient-reported symptom scores for antibiotic-naive patients at diagnosis, 6 months, and 1 year. Symptoms are reported as absent to severe (0-1). Each column represents a patient at a given time point. **(E)** Boxplot of average symptom score over time in antibiotic-naive patients. Each line is a patient. The average symptom score is the average of all patient-reported symptoms. **(F)** Bubble plot showing Pearson correlation coefficients of symptom and clinical values in antibiotic-naive patients at diagnosis. Size and color indicate the magnitude and direction of correlation respectively. Correlations were computed with the rcorr function from the SVA package in R.

**Table 1:** Characteristics of the longitudinal patient cohort recruited in this study. Key demographic and clinical features of the cohort are summarized.

|  | All Patients | Antibiotic Naive Patients | Controls |
| --- | --- | --- | --- |
| **Demographics** |  |  |  |
| Diagnosis | 49 | 21 | 50 |
| 2-3 Weeks | 49 | 21 | - |
| 6 Months | 47 | 20 | 45 |
| 1 Year | 46 | 20 | 46 |
| Age<sup>1</sup> | 50.1±15.4 [18, 73] | 52.3±10.9 [26, 69] | 51.8±15.1 [23, 76] |
| Female | 22 | 10 | 22 |
| Male | 27 | 11 | 28 |
| **Serology (Positive, Negative, Equivocal, Not Done)** |  |  |  |
| Diagnosis | (41,8,0,0) | (17,4,0,0) | - |
| 2-3 Weeks | (45,3,0,1) | (19,2,0,0) | - |
| 6 Months | (13,5,2,27) | (5,2,1,12) | - |
| 1 Year | (22,22,1,1) | (10,10,0,0) | - |
| **Coinfection** |  |  |  |
| None | 42 | 17 | - |
| Babesia | 4 | 2 | - |
| Powassan | 1 | 0 | - |
| Not Done | 2 | 2 | - |
| **Erythema Migrans** |  |  |  |
| None at Baseline | 14 | 2 | - |
| single | 24 | 11 | - |
| 2-3 | 3 | 2 | - |
| 4+ | 8 | 6 | - |
| Largest Dimension (cm)<sup>1</sup> | 13.3±7.8 [5, 45] | 14.8±9.3 [8, 45] | - |
| **Antibiotics** |  |  |  |
| Doxycycline | 37 | 13 | - |
| Amoxicilin | 10 | 7 | - |
| Ceftriaxone | 5 | 0 | - |
| Cefuroxine | 1 | 0 | - |
| Augmentin | 2 | 1 | - |
| <sup>1</sup>Summary statistics reported as: Mean ± SD [Range] |  |  |  |

All patients presented with physician-diagnosed EM and were confirmed for seropositivity by the C6 ELISA, which measures host antibodies to the conserved C6 peptide of the *Borrelia* VlsE protein. Most antibiotic-naïve patients produced C6 antibodies at T1 (81%) and T2 (90%), with approximately half of the cohort maintaining antibody levels up to one year post-infection **(Fig. 1c**, **Table 1)**. At T1, T3, and T4, patients reported the severity of 12 symptoms associated with LD using an 8 cm visual analogue scale. Symptoms were reviewed to determine whether they were attributable to LD or a confounding condition. General health data, including body mass index (BMI), temperature, blood pressure, and pulse rate, were also collected.

During the acute disease, fatigue was the most frequently reported symptom (52%), followed by joint pain (38.1%), headache (33.3%), and stiff neck (33.3%) **(Fig. 1d)**. Although symptom presentation varied across the cohort—from individuals reporting multiple symptoms to a small subset presenting solely with an EM lesion and C6 seropositivity—most patients experienced their peak symptom burden at the time of diagnosis, as reflected by the elevated average symptom score at T1 **(Fig. 1e)**. Following antibiotic treatment, symptoms resolved in most patients, with only a few reporting mild symptoms attributable to LD at 6 months or at 1 year post-treatment **(Fig. 1d-e)**. At diagnosis, patient-reported symptoms were highly correlated with each other, except tingling/abnormal sensation, which showed no positive correlation with other symptoms **(Fig. 1f)**. BMI also correlated strongly with diastolic blood pressure, consistent with previous findings^23^.

Although patients often could not identify the date of tick bite to establish the onset of infection, the diameter of the EM rash serves as a rough indicator of disease duration, as the rash expands radially over time. Additionally, the presence of multiple EM lesions at secondary sites has been used as a marker of disseminated infection in prior studies^24,25^. While we observed minimal correlations between symptom severity and EM dimensions, the number of EM lesions positively correlated with fatigue (r=0.58) and stiff neck (r=0.46), suggesting that disseminated disease may mildly exacerbate symptomatology **(Fig. 1f)**. Symptom presentation was largely independent of age, with the exception that older patients reported fewer concentration and memory problems. Together, these data demonstrate the heterogeneous nature of LD symptoms during the acute phase and their resolution following antibiotic treatment, highlighting variability in the magnitude and nature of clinical manifestations among different individuals.

### Temporal dynamics of circulating immune proteins in acute LD

To understand the acute immune response to LD and whether it is reflected in changes in circulating mediators, we measured the relative levels of 182 circulating immunomodulatory proteins in plasma samples from patients and controls using Olink Inflammation and Immune Response panels. In antibiotic-naive patients, 105 proteins were significantly induced at diagnosis compared to controls. This number decreased sharply following antibiotic treatment, and by T2 only the peroxiredoxin family member PRDX5 (an antioxidant that protects against inflammatory stress)^26^ remained significantly elevated. By 6 months and 1 year post-treatment, no statistically significant differences were observed between patients and controls **(Fig. S1b)**. Comparisons of plasma protein levels between patients who were already on antibiotics at study entry, controls, and antibiotic-naive patients at T1 revealed that even a short prior period of antibiotic treatment reduces circulating immune mediators **(Fig. S1c)**.

Human subjects exhibit significant baseline immune variation, necessitating careful analysis to discern true biological signals from inter-individual noise **(Fig. S1d)**. To account for this variability, we conducted pairwise differential abundance analysis between different time points for each patient. Clustering the resulting log fold-changes revealed fast- and slow-resolving immune protein signatures that were significantly induced at T1 but resolved at different rates over time **(Fig. 2a and S1e)**. The fast-resolving cluster primarily included inflammatory cytokines, chemokines, and growth factors. The most highly induced were the IFN-γ-inducible chemokines CXCL9, CXCL10, and CXCL11, as well as certain cytokines such as IL-6, IL-10, IFN-γ, CCL19, IL-17C, CCL3 (MIP1α), CXCL8 (IL-8), and CCL4 (MIP1β), along with enzymes like SIRT2. These proteins, mostly cytokines and chemokines, are known to act rapidly in mediating immune responses and are regulated by transient transcriptional programs and rapid clearance mechanisms^27,28^. Consequently, they peaked at T1, resolved rapidly to sub-baseline levels by T2, and returned to baseline levels observed at T3 and T4. Pathway analysis showed that these proteins mapped predominantly to immune cell chemotaxis, leukocyte migration, and G protein-coupled receptor signaling **(Fig. 2b)**.

**Figure 2:**
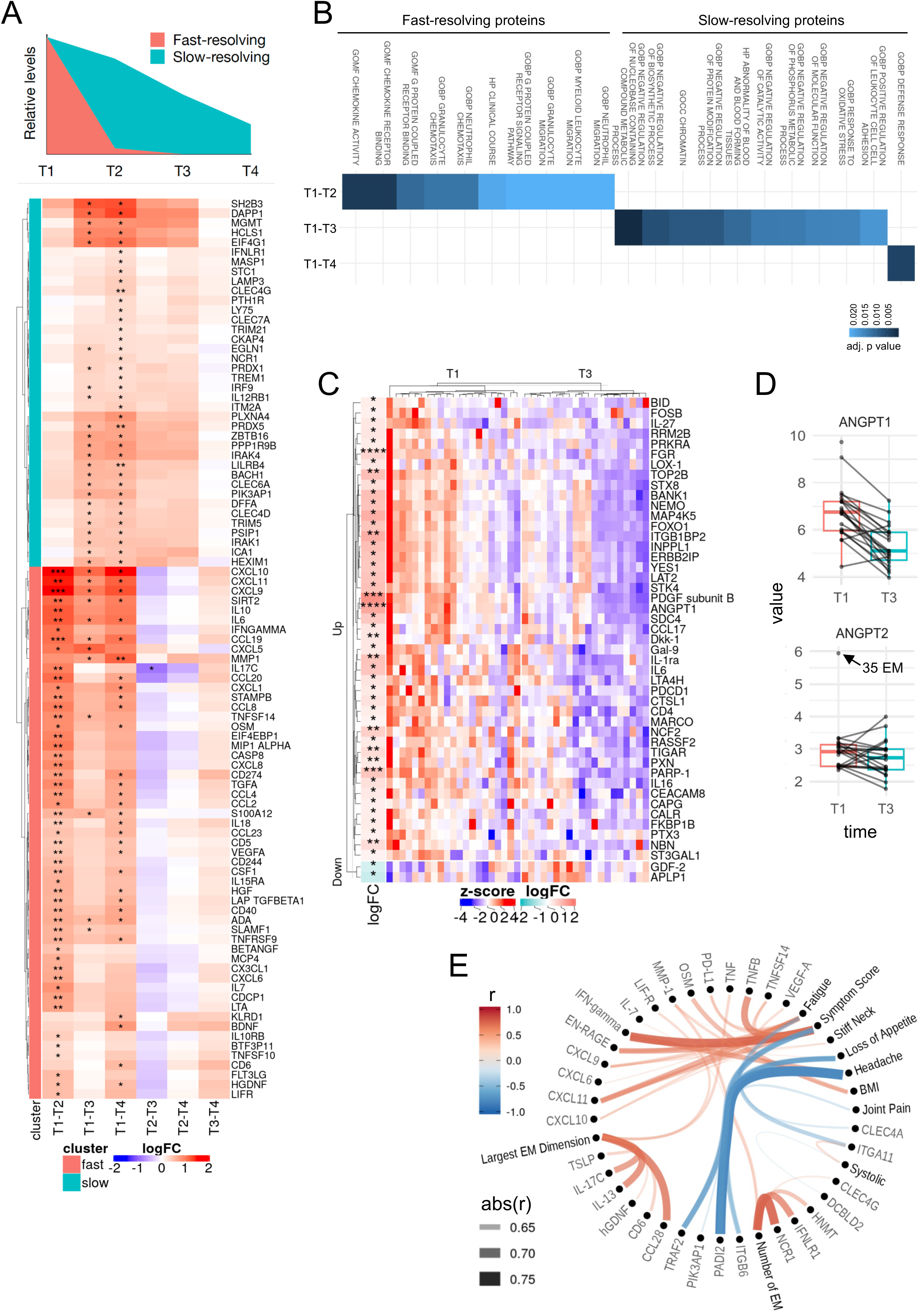
Differential expression of circulating proteins and their correlation with symptoms in acute disease. **(A)** Cartoon trajectories (top) and heatmap (bottom) of fast- and slow-resolving proteins, illustrating their distinct temporal dynamics in plasma. The heatmap shows the differential expression of proteins from the Olink Immune Response and Inflammation panels, tested across pairs of time points (indicated at the bottom) in antibiotic-naive patients. Statistical significance is marked by asterisks in each cell (*p < 0.05, **p < 0.01, ***p < 0.001). Log fold change (logFC) values were hierarchically clustered using the Ward.D2 method (hclust in R). The resulting tree was cut to reveal two protein clusters: fast- and slow-resolving. **(B)** Heatmap showing differentially induced pathways at the indicated time points. p-values obtained from GSEA with the clusterProfiler R package on the Hallmark, C2, C5, and C7 gene sets from MSigDB are shown. GSEA was run on differentially expressed proteins discovered during the indicated pairwise comparisons. **(C)** Heatmap showing z-score normalized protein expression from the Olink Cardiovascular II, Metabolism, and Organ Damage panels (rows) in antibiotic-naive patients (columns). Log fold change values and significance for each protein, compared pairwise between T1 and T3, are shown on the left. Differential expression was computed using the R package limma. *p<0.05, **p<0.01, **p<0.001. **(D)** Box plots showing the relative levels of the ANGPT proteins over time. Each line represents a patient. Unpaired individual at T1 with highest ANGPT expression represents a patient with 35 EM. **(E)** Hierarchical edge bundling plot of significant Pearson correlations (p<0.05) between circulating proteins and patient-reported symptom scores in antibiotic-naive patients at diagnosis. Proteins and symptoms with similar correlation patterns are grouped together by hierarchical clustering. Line width and color indicate the strength and direction of the correlation respectively. Correlations were computed using the rcorr function from the sva R package.

In contrast, the slow-resolving cluster contained many inhibitory proteins associated with immune regulation and tissue homeostasis. Notable proteins included SH2B3 (an inhibitor of cytokine signaling via JAK-STAT pathways)^29^, STC1 (a pleiotropic glycoprotein with anti-inflammatory properties)^30-32^, CLEC4G (which inhibits T cell activation and antigen presentation)^33,34^, EGLN1 (which reduces HIF-1α-driven inflammation)^35^, PRDX1 and PRDX5 (antioxidants that lower reactive oxygen species and attenuate inflammatory signaling)^26^, PLXNA4 (which regulates immune cell trafficking)^36^, LILRB4 (an inhibitory receptor on myeloid cells that downregulates inflammatory cytokine production)^37^, BACH1 (a heme-regulated transcriptional repressor involved in oxidative stress and immune regulation)^38^, and HEXIM1 (a regulator that represses RNA polymerase II activity, modulating inflammatory gene expression)^39,40^. These proteins decreased gradually after peaking at T1 and persisted at elevated levels until T3. Certain proteins, such as CLEC4G and STC1, did not significantly decline until T4, suggesting that while most slow-resolving proteins resolved by T3, a subset exhibited prolonged persistence **(Fig. 2a)**. Pathway analysis mapped these proteins predominantly to processes related to negative regulation of molecular function, catalytic activity, and metabolic processes **(Fig. 2b)**. Together, these findings reveal distinct temporal patterns of immune modulation, with fast-resolving proteins mediating acute inflammation and slow-resolving proteins potentially contributing to a sustained regulatory environment during the later stages of acute infection.

### Vascular and symptom-associated protein signatures

To investigate the broader physiological effects of LD, we measured an additional 276 proteins related to metabolism, cardiovascular health, and organ damage using Olink proteomics panels. Pairwise differential abundance analysis revealed that 45 proteins were upregulated and 2 were downregulated at T1 relative to T3 **(Fig. 2c)**. Notably, vasoprotective proteins such as angiopoietin-1 (ANGPT1) and platelet-derived growth factor-B (PDGFB), which promote endothelial stability, were significantly elevated at diagnosis. ANGPT1 and PDGFB signal in endothelial cells and are essential for maintaining vascular homeostasis^41,42^. ANGPT1 functions as an agonist of the Tie2 receptor, enhancing vascular stability and integrity, while ANGPT2 acts as its competitive antagonist, destabilizing endothelial barriers and exacerbating vascular dysfunction in inflammatory conditions. ANGPT2 levels remained stable in most patients but were elevated in one individual with severe disease **(Fig. 2d)**. Additionally, growth differentiation factor 2 (GDF-2), a secreted ligand of the TGF-β superfamily and a vasodisruptive protein that inhibits angiogenesis and endothelial cell growth^43^, was significantly downregulated at T1 compared to T3. These findings highlight the balance between endothelial-protective and disruptive responses in LD and the potential role of ANGPT1-Tie2 signaling in modulating vascular homeostasis during acute disease.

We next examined the relationship between circulating proteins and clinical manifestations by correlating protein levels at T1 with symptom severity (average of all patient-reported symptom scores), individual symptoms, and clinical parameters, focusing on analytes from the immune response and inflammation panels. Symptom severity and fatigue were strongly associated with two distinct protein modules. The first module included IFN-γ and IFN-γ-inducible proteins such as CXCL9, CXCL10, CXCL11, PD-L1, and the antimicrobial peptide EN-RAGE. The second module comprised TNF family members TNFSF14 and TNF-β, as well as the vascular growth factor VEGF-A **(Fig. 2e)**. These findings confirm the presence of an IFN-γ-associated Th1 cytokine signature in LD^21,44-47^, linking it to symptom severity. Dermatological responses, such as EM dimension, correlated with a distinct type 2 cytokine signature (IL-13, TSLP, and CCL28) typically associated with allergic or parasitic responses^48,49^, as well as the pro-inflammatory cytokine IL-17C, predominantly produced by keratinocytes. While the type 2 cytokines may help modulate inflammation and promote tissue repair, IL-17C likely enhances local immunity by recruiting immune cells and strengthening barrier defenses^50^. These cytokines likely represent localized efforts to balance immune activation and tissue repair, with the immunomodulatory properties of tick saliva^51^ potentially influencing these dynamics to facilitate pathogen establishment while simultaneously triggering repair mechanisms. Conversely, the number of EM lesions, an indicator of disseminated disease, correlated with the epithelium-restricted IFNLR1 and the triggering receptor for NK cell cytotoxicity NCR1 (NKp46), whose functions in plasma remain unclear **(Fig. 2e)**. Together, these data reveal distinct IFN-γ-associated and type 2 cytokine/IL-17C signaling programs contributing to symptom severity and dermatological responses in LD.

Building on these associations, we applied LASSO (least absolute shrinkage and selection operator) regression models to determine if circulating protein signatures could classify patient samples from controls at different time points. LASSO regression was specifically chosen to prevent overfitting, given the high dimensionality of our dataset. Models trained on the immune protein data achieved moderate accuracy at T1 and T2 (AUC = 0.684 and 0.61, respectively) but lost predictive power by T3 and T4 **(Fig. S1f)**. These models, validated using nested Monte Carlo cross-validation, provide conservative performance estimates and highlight the diagnostic potential of circulating immune protein signatures early in infection. However, their limited accuracy underscores the need for larger cohorts or the integration of additional omics layers to improve performance.

Rare clinical features were explored using principal component analysis (PCA) of protein signatures, which revealed unique outliers among patients. Outlier patients presented with fever and the highest levels of IFN-γ-inducible and other inflammatory cytokines from the fast-resolving cluster, which were associated with symptom severity at T1 **(Fig. S2a)**. Patients with multiple EM lesions did not exhibit distinct protein signatures, suggesting that disseminated disease does not necessarily alter the plasma proteome **(Fig. S2b)**. Similarly, two patients with *Babesia* co-infection showed no notable differences from other LD patients **(Fig. S2c)**. These results suggest that among clinical parameters, fever is associated with the largest shifts in circulating inflammatory proteins, highlighting its impact on systemic immune signatures in LD.

### Metabolite-protein associations reveal a balance of endothelial protection and dysfunction in acute LD

We next analyzed 1244 metabolites in patient plasma using Metabolon’s platform to explore the metabolic changes underlying acute LD. An inter-omic community analysis of proteomics (Olink Inflammation, Immune Response, Cardiovascular, Organ Damage, and Metabolic panels) and metabolomics data allowed us to identify physiologically related analytes. Among the three largest communities with analytes predominantly upregulated at T1 (Community 1 with 29 members, Community 3 with 26 members, and Community 4 with 22 members; **Fig. S3a**), Communities 4 and 1 emerged as key signatures representing endothelial disruption and protection, respectively **(Fig. 3a)**.

**Figure 3:**
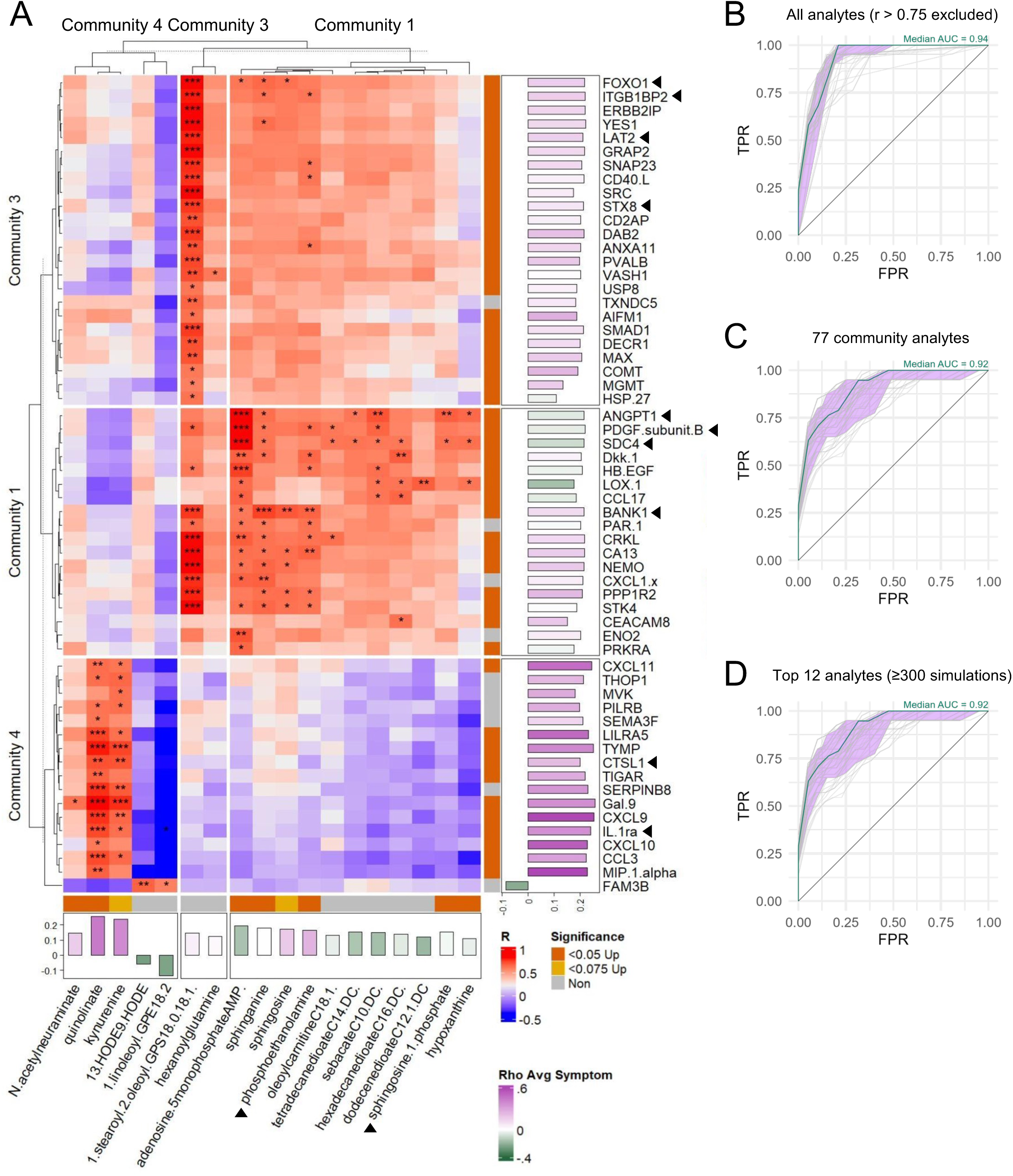
Integrated community analysis and diagnostic modeling of circulating proteins and metabolites in LD. **(A)** Heatmap showing correlations between proteins (rows) and metabolites (columns) within the three largest communities identified by interomic community analysis. Cell colors indicate the strength and direction of the protein-metabolite correlation; the correlation was statistically significant (FDR-corrected) for cells containing an asterisk. Features within each community were clustered by hierarchical clustering based on the correlation values. Annotations immediately beside the heatmap indicate whether the protein or metabolite was significantly upregulated at T1 compared to T3. Floating bars adjacent to the heatmap represent each feature’s contribution to PC1 of its respective community, with bar height reflecting the strength and direction of the contribution (positive or negative); longer bars indicate stronger influence. Bar colors denote Spearman correlations between symptom scores and expression levels: green for negative correlations and purple for positive correlations. **(B, C, D)** ROC curves from LASSO regression models with Monte Carlo cross-validation, classifying patients at T1 and T3 using data from all Olink panels and Metabolon analytes measured in this study. Each plot shows individual ROC curves from 500 simulations (light gray lines), the median ROC curve (green), and the 95% confidence interval (purple ribbon). **(B)** includes analytes with pairwise correlations below 0.75 and those from the top three communities. **(C)** uses analytes from the top three communities only. **(D)** uses a reduced set of 12 key analytes consistently selected in ≥ 300 of the 500 simulations. The 12 analytes used in (D) are indicated by a black triangle next to the analyte name in (A).

Community 4 included IFN-γ-inducible chemokines (CXCL9, CXCL10, CXCL11), MIP-1α, anti-angiogenic factors (e.g., GAL-9, TIGAR, SEMA3F, THOP1, CTSL1)^52-56^, and proteins involved in vascular inflammation and thrombosis (e.g., TYMP, LILRA5, PILRB, IL-1RA)^57-61^. Metabolites of the tryptophan metabolism pathway, including kynurenine and quinolinate, were also prominent. IFN-γ induces the expression of indoleamine 2,3-dioxygenase 1 (IDO1), which catalyzes the rate-limiting step in tryptophan metabolism, resulting in bioactive metabolites that affect the immune, nervous, and vascular systems^62^. Quinolinic acid, in particular, has been implicated in adverse outcomes such as post-treatment and neuropsychiatric complications in LD^63,64^. Features from Community 4 were strongly associated with disease severity, underscoring the IFN-γ-IDO1-quinolinate axis as a key driver of endothelial disruption and inflammation during acute disease **(Fig. 3a)**.

Conversely, Community 1 contained proteins supporting endothelial protection, including ANGPT1, PDGFB, syndecan-4 (SDC4), and heparin-binding EGF-like growth factor (HBEGF)^42,65,66^. It also featured intermediates of sphingolipid metabolism, such as sphingosine, sphingosine-1-phosphate (S1P), phosphatidylethanolamine, and sphinganine, which regulate endothelial barrier integrity^67^. Additionally, this community included regulators of immune cell adhesion, endothelial stress, and angiogenesis such as CCL17, CEACAM8, CRKL, DKK1, LOX1, PRKRA, and STK4^68-74^. Among these, S1P is particularly important for immune cell trafficking and vascular integrity and has been shown to inversely correlate with disease severity in inflammatory conditions like COVID-19^75^. Together, Communities 4 and 1 represent opposing endothelial disruption and protection respectively, reflecting the intricate balance of vascular responses during LD. Community 3 contained several immunoregulatory proteins, some involved in vascular remodeling, but showed less clear associations with metabolites or disease severity.

To evaluate the diagnostic utility of these findings, we applied LASSO regression models to integrated plasma proteomics and metabolomics data, excluding highly correlated analytes. This comprehensive model demonstrated high accuracy (AUC = 0.94) in distinguishing patients at T1 (diagnosis) from those at T3 (6 months; **Fig. 3b**). A reduced model using only the 77 analytes identified in our community analysis retained a high level of discriminatory power (AUC = 0.92), suggesting that key biological pathways — particularly those involved in inflammatory and vascular-protective responses — are critical for distinguishing acute LD from later time points (**Fig. 3c**). These models were built using 500 iterations of Monte Carlo cross-validation, ensuring robustness by repeatedly fitting the model on random subsets of the data. Further refining the predictors to the 12 community analytes that appeared in at least 300 of the simulations did not affect the model’s accuracy (AUC = 0.92; **Fig. 3d**). These findings highlight the potential for developing diagnostic tools that leverage integrated multiomics data. However, additional studies are needed to evaluate the specificity of these tools for LD relative to other infectious and inflammatory conditions.

In an orthogonal approach to the community analysis, pathway analysis of metabolomics data confirmed the upregulation of tryptophan and sphingolipid metabolism **(Fig. 4a)**. Additional upregulated modules were identified, including beta-oxidation, a process that breaks down fatty acids for energy production under low glucose or high energy demand^76^. Similar metabolic alterations— including increased beta-oxidation and sphingolipid pathway activity—have also been reported in independent studies of early LD^77,78^, supporting the robustness of these signatures across cohorts and analytical platforms. The main product of beta-oxidation, acetyl-CoA, is funneled into ketogenesis, producing ketone bodies such as 3-hydroxybutyrate (BHBA), acetoacetate, and acetone. Metabolites mapping to these pathways were significantly elevated at T1 compared to later time points **(Fig. 4b-d)**. Quinolinate levels peaked at T1, accompanied by a subtle decrease in plasma tryptophan levels, reflecting increased flux through the kynurenine pathway **(Fig. 4b)**. Increased ketogenesis products, such as BHBA and acetoacetate, suggest that patients may be experiencing energetic stress, yet these ketones, like S1P, also have vasoprotective properties that support endothelial homeostasis^79,80^.

**Figure 4:**
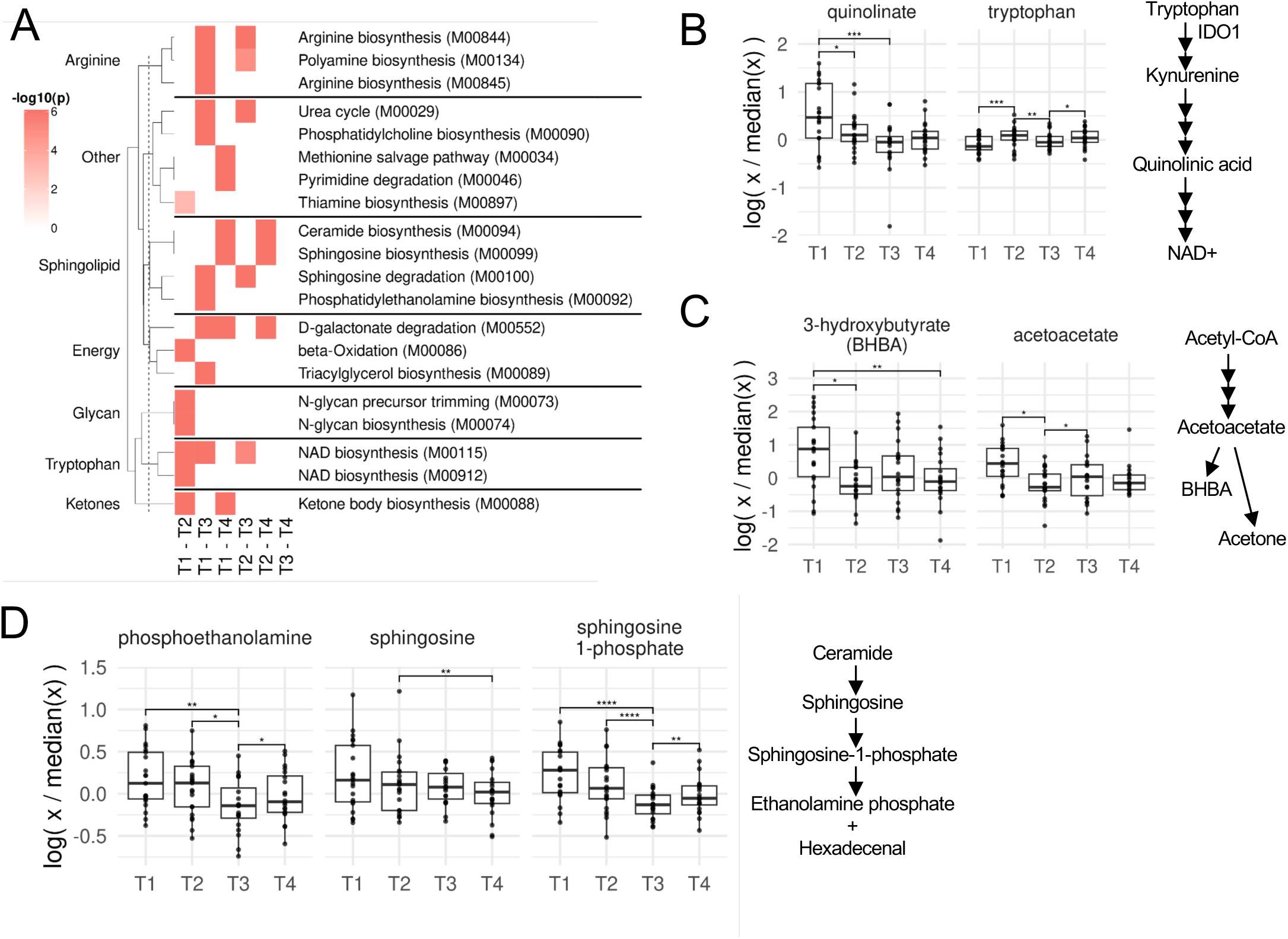
Metabolic pathway enrichment and metabolite abundance in key pathways over time. **(A)** Heatmap showing enriched metabolic pathway modules from KEGG. Enrichment was performed using the R package FELLA, with significantly differentially expressed metabolites supplied as input. Differential expression comparisons were made between the indicated time points. **(B, C, D)** Box plots of normalized metabolite abundance across time points. p-values indicate significant temporal variation in metabolite abundance, as determined by a likelihood ratio test comparing models with and without time as a factor.

Quinolinate correlated significantly with pro-inflammatory proteins (e.g., MIP-1α, CXCL9, CXCL10, and CXCL11) and symptom severity **(Fig. S3b)**, reinforcing its role in the IFN-γ-IDO1-quinolinate axis. Conversely, BHBA correlated significantly with palmitoleate (16:1n7), a lipokine known to regulate beta-oxidation via PPAR-α^81^ and reduce inflammation through multiple mechanisms^82^. Both BHBA and S1P correlated strongly with ANGPT1 and PDGFB, suggesting a shared link to vasoprotective signaling and implicating these metabolites in endothelial barrier protection during acute disease. These findings confirm the induction of opposing endothelial-protective and inflammatory plasma signatures, emphasizing the dynamic interplay between vascular stress and homeostasis in acute LD.

### Minimal immune activation in PBMCs except in severe disseminated disease

We investigated changes in the abundance and activation of peripheral immune cells during acute LD to assess systemic immune engagement. Immunophenotyping of PBMCs from all subjects in our cohort was performed using four multicolor flow cytometry panels, profiling T cells, B cells, monocytes, dendritic cells (DCs) and natural killer (NK) cells **(Table S1)**. These panels were analyzed by a pipeline developed in-house based on unbiased clustering followed by manual cluster merging and annotation to identify 41 distinct immune cell populations **(Fig. S4-S5 and Table S2)**. PCA of z-score normalized cell population abundances revealed that, like controls, patient samples clustered by identity rather than time-point, except for one individual with severe disseminated disease who presented with 35 distinct EM lesions and flu-like symptoms. This clustering pattern indicates that the impact of infection on peripheral immune cell composition was smaller than preexisting inter-individual variability (**Fig. 5a**). Differential abundance analysis conducted within each patient over time revealed consistently elevated plasmablasts at T1 **(Fig. 5b)**, consistent with antibody production against the C6 peptide of the *B. burgdorferi* VlsE protein **(Fig. 1c)**. Other immune cell populations exhibited minimal or inconsistent changes across time points **(Fig. S6a)**. Associations of cell types with symptoms **(Fig. S6b)** and circulating proteins **(Fig. S6c)** were sparse and appeared driven by one outlier patient with 35 EM lesions, likely representing artifacts of multiple testing rather than true biological signals. These results indicate that acute LD does not induce broad changes in PBMC composition, with the exception of a modest humoral response.

**Figure 5:**
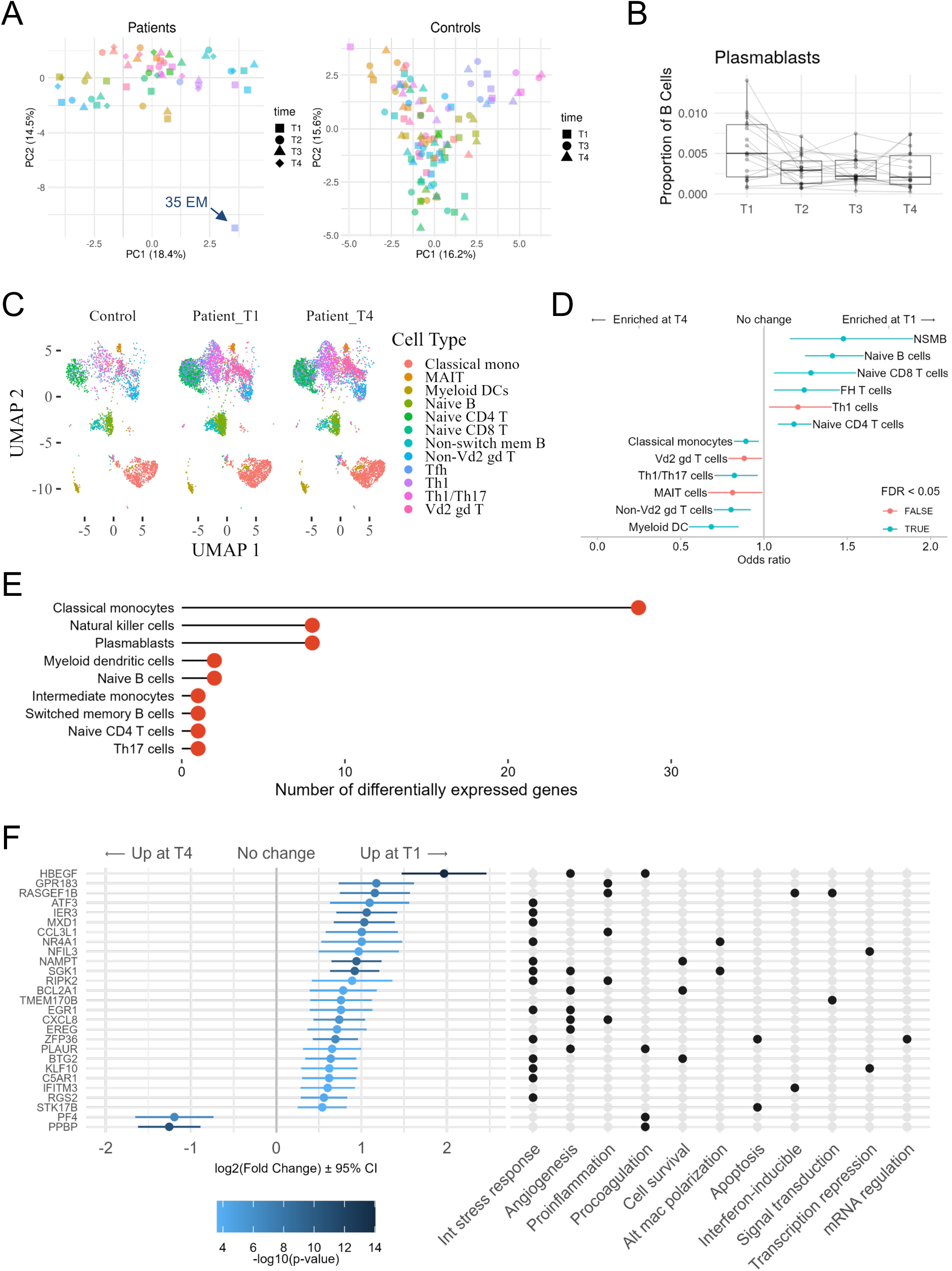
Minimal peripheral changes in acute LD. **(A)** PCA of PBMC abundances from flow cytometry, shown separately for patients (left) and controls (right). Abundances were normalized across all patient samples at each time point. Point color denotes subject identity, and shape denotes time point. **(B)** Paired boxplots showing changes in plasmablasts as a proportion of total B cells in patients over time. **(C)** UMAPs of PBMCs from patients and controls. Shown are aggregated scRNA-Seq profiles from two patients with matched T1 and T4 samples and four healthy controls, displaying only cell types with significant or trending changes in abundance over time. For visualization only, equal numbers of cells were randomly subsampled from each condition to ensure comparable representation across groups. **(D)** Forest plot of cell types with significantly different abundance between T1 and T4 in PBMC scRNA-Seq data. Differential abundance was assessed using a generalized linear model with time as the main predictor, adjusting for subject ID. Horizontal bars indicate confidence intervals around odds ratios. An odds ratio of one suggests no difference between groups. **(E)** Plot showing the number of differentially expressed genes per cell type in PBMC scRNA-Seq. **(F)** Left: Forest plot showing the log2 fold change in gene expression (T1 vs. T4) for classical monocytes, with error bars representing 95% confidence intervals and the colors indicating significance (−log10 p-value). Right: The same genes mapped to functional categories^73^, with each column representing a function and filled circles denoting gene inclusion in a specific category. Both plots are aligned by gene rows.

To investigate transcriptional changes in PBMCs and validate flow cytometry findings, we performed single-cell RNA-Seq (scRNA-Seq) on a subset of samples collected at T1 and T4. Among 27 identified cell types **(Fig. S7a)**, naïve B cells, naïve CD4+ and CD8+ T cells, follicular helper T cells (Tfh), and non-switched memory B cells (NSMB) exhibited small increases (odds ratio ≤ 1.5) at T1 compared to T4 **(Fig. 5c-d)**. Minor fluctuations in Th1 cells did not remain significant after multiple testing corrections, and no robust increases in effector T or B cells were observed (**Fig. 5d and S7b**). Differential gene expression analysis identified 27 differentially expressed genes (DEGs) in classical monocytes, 8 in NK cells, and 4 in plasmablasts, with few or no DEGs in other cell types (**Fig. 5e and Table S3**). Classical monocytes displayed an unconventional transcriptional program characterized by endothelial-protective and patrolling signatures rather than inflammatory profiles. Upregulated genes in these cells at T1 included those involved in angiogenesis, wound healing, dissolution of blood clots, and endothelial survival (e.g., *HBEGF, PLAUR, EREG, EGR1, SGK1*), while genes associated with platelet activation and clot formation (*PF4, PPBP*) were downregulated **(Fig. 5f and S8)**. Additional upregulated genes were linked to the integrated stress response (e.g., *ATF3, SGK1, NR4A1, IER3, EGR1, MXD1, C5AR1, NAMPT, BTG2, RGS2*), transcriptional repression (e.g., *NFIL3, KFL10*), survival and growth regulation (e.g., *BCL2A1, NAMPT, BTG2*), alternative macrophage polarization (e.g., *NR4A1, SGK1*), and inflammation-limiting RNA-binding proteins such as *ZFP36*, which attenuates pro-inflammatory signaling by destabilizing cytokine mRNAs^73^. Pro-inflammatory genes like *IL-8, CCL3L1, RIPK2,* and *IFITM3* were also upregulated, although they did not dominate the expression profile **(Fig. 5f)**. These findings highlight an endothelial-protective role for classical monocytes during acute infection, indicating a non-canonical function for these typically inflammatory cells, while the overall PBMC transcriptional response remained largely quiescent.

Despite the limited peripheral immune activation observed across most patients, one individual with severe disseminated disease and 35 EM lesions was a clear outlier in PCA at T1 **(Fig. 5a)**. This patient exhibited markedly elevated circulating cytokines and chemokines, including IFN-γ-inducible chemokines (CXCL9, CXCL10, CXCL11) and other inflammatory mediators **(Fig. 6a, Table S4)**. Proteins associated with the release of Weibel-Palade bodies (WPBs) from endothelial cells, such as ANGPT2 and IL-8^83^, were substantially elevated compared to other patients, reflecting significant disruption of vascular homeostasis **(Fig. 6a, 6b, 2d, and Table S5)**. Plasma metabolite analysis showed increased levels of quinolinate, inosine, and sphingolipids, alongside reduced tryptophan, indicating heightened immune-metabolic activity that was similar in nature but substantially greater in magnitude compared to other patients **(Fig. 6c, Table S6)**. Immune profiling revealed a marked expansion of effector T cells and an atypical population of monocytes co-expressing high levels of both CD14 and CD16 (CD14++CD16++), exceeding the canonical intermediate monocyte subset **(Fig. 6d)**. The CD14++CD16++ subset expressed high levels of TNF-α, CD40, and CD86 but reduced HLA-DR, suggesting a hyperinflammatory state with potential immunoparalysis^84^ **(Fig. 6e)**. Notably, no similar immune cell activation was observed in other patients, including those with up to 27 EM lesions **(Fig. S9)**. These findings demonstrate that while PBMC responses in acute LD are generally quiescent, systemic immune activation can occur in certain individuals with severe disseminated disease marked by exaggerated cytokine production, metabolic shifts, and vascular dysfunction.

**Figure 6:**
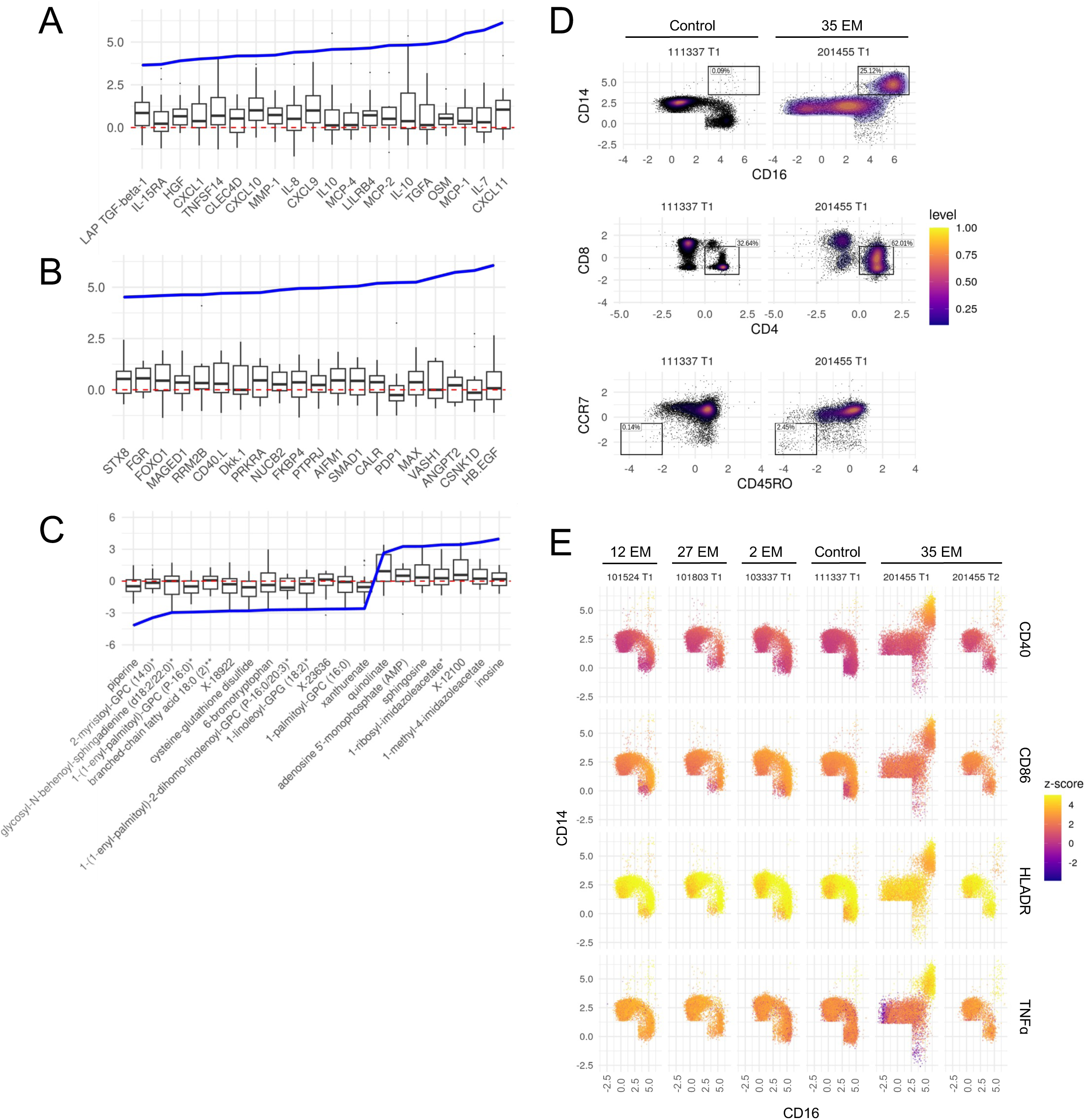
Dramatic changes in circulating mediators and immune cell abundance in a case with severe disseminated disease. **(A, B, C)** Olink plasma proteins (A, B) and metabolite (C) z-scores for an outlier patient with 35 EM at diagnosis (blue line). Boxplots show z-scores of patient samples. Z-scores were computed on all patient and control samples at all time points. Plots display results from the Olink Inflammation and Immune response panels (A), Olink Metabolism, Cardiovascular II, and Organ Damage panels (B), and metabolomic data (C). Only the top 20 differentially regulated mediators are shown. **(D)** Flow cytometry plots of monocytes and T cells from an outlier patient at diagnosis (201455 T1), and a representative age/sex-matched healthy control (111337 T1). Monocytes were defined as CD14+Dump- and T cells were defined as CD3+ (parent gates not shown). Level indicates relative cell density. **(E)** Representative flow cytometry plots of monocyte activation for patients with varying numbers of EM at diagnosis, and a healthy control. Color indicates z-score for the marker (CD40, CD86, HLADR or TNF-α) indicated on the right. Z-scores were computed across all patient and control cells at all time points.

### Skin resident cells predominantly shape circulating immune and metabolic signatures in LD

To validate our PBMC findings and investigate immune responses at the site of infection, we analyzed publicly available bulk transcriptome data from PBMCs and EM skin lesions of LD patients **(Table 2)**. Immune cell enrichment analysis using xCell^85^ revealed modest changes in immune cell composition in PBMCs at diagnosis relative to controls, including enrichment of monocytes and plasma cells— findings broadly consistent with our flow cytometry data. In contrast, EM lesions exhibited robust immune cell infiltration, with marked enrichment of monocytes, macrophages (particularly M1), neutrophils, plasmacytoid dendritic cells (pDCs), basophils, mast cells, and NK cells **(Fig. S10a)**. These results support the limited peripheral immune activation observed in our cohort and highlight the site-specific nature of the immune response in acute LD.

**Table 2:** Characteristics of the public gene expression datasets analyzed in this study. Gene expression data was downloaded from the Gene Expression Omnibus (GEO) and analyzed as described in the methods.

| Title | Type | Tissue | Conditions | GEO Accession |
| --- | --- | --- | --- | --- |
| Identification of a Molecular Signature for Acute Lyme Disease by Human Transcriptome Profiling | RNAseq | PBMC | Patients (n=29), Controls (n=13)<br>0, 21, 182 days | GSE63085 |
| Transcriptome Assessment of Erythema Migrans Skin Lesions | Microarray | Skin | Patients (n=18), Controls (n=27)<br>EM, healthy skin | GSE154916 |
| Single Cell Immunophenotyping of Lyme Erythema Migrans | scRNAseq | Skin | Patients (n=10)<br>EM, autologous healthy skin | GSE169440 |

Given the minimal activation of immune cells in the periphery, we sought to identify the source of cytokines, chemokines, and other circulating proteins detected in plasma during acute disease. To address this, we reanalyzed public scRNA-Seq data from skin punch biopsies^86^ **(Fig. S11-S13, Table 2)**, which revealed a significant increase in immune cell abundance in EM lesions compared to autologous healthy skin **(Fig. 7a-b and S10b)**. Surprisingly, the dominant cytokine signature detected in patient plasma—including IFN-γ-inducible chemokines (CXCL9, CXCL10, CXCL11), IL-8, MMP-1, CCL19, CXCL1, and CX3CL1, as well as inhibitory proteins like PD-L1 and LILRB4^37,87,88^—was primarily upregulated in skin-resident cells. These included fibroblasts, keratinocytes, endothelial cells (lining the interior surface of all blood vessels and regulating vascular permeability), vascular smooth muscle cells (vSMCs; contractile cells surrounding large blood vessels that regulate vascular tone and blood pressure), and pericytes (supportive cells encasing capillaries and small vessels that maintain vascular stability and mediate blood flow) **(Fig. 7c)**. A unique population of skin-infiltrating T cells (T cell1) was identified as the primary source of IFN-γ **(Fig. 7c and S12c)**. In contrast, infiltrating myeloid cells such as macrophages and CD1c+ DCs showed minimal cytokine production except for IL-18, while upregulating the immunoinhibitory gene LILRB4. These results link the circulating immune signature to the site of infection, suggesting that skin-resident cells are key contributors to the major cytokine and chemokine response in LD.

**Figure 7:**
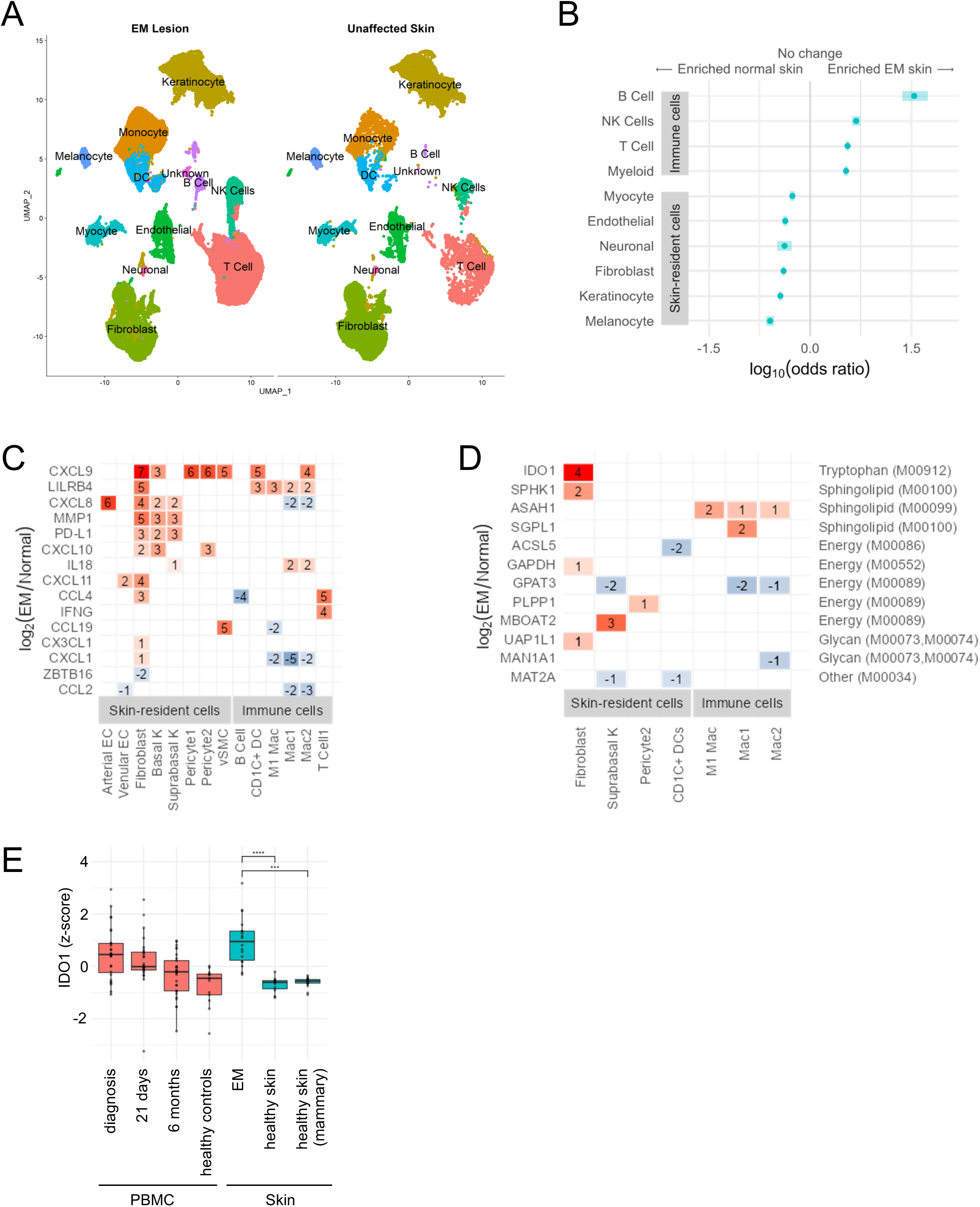
Skin immune responses reflect plasma protein and metabolic signatures. **(A)** UMAPs of scRNA-Seq profiles from matched EM lesion and unaffected autologous skin biopsies (GSE169440), annotated by major cell types. Immune cell populations—including T cells, NK cells, B cells, dendritic cells, and monocytes—are expanded in EM lesions relative to unaffected skin, highlighting localized immune infiltration at the site of infection. **(B)** Forest plot of cell types with significantly different abundance in normal vs. EM skin, based on reanalysis of the same scRNA-Seq dataset. Differential abundance was calculated using a generalized linear model that included subject ID and skin condition (normal or EM). Horizontal bars represent confidence intervals around odds ratios for each cell type. A log10(odds ratio) of zero indicates no difference between the groups. **(C, D)** Immune genes (C) and enzyme genes (D) with significant differential expression in skin, after controlling for the false discovery rate (FDR). Cell identities, grouped into skin-resident cells and immune cells, are shown along the x-axis. Tile colors and numbers represent log2 fold changes in EM versus normal skin, with white indicating non-significant changes. In (D), the y-axis on the right specifies the KEGG module (referenced in Fig. 4a) where each enzyme gene is involved. **(E)** Box plots showing z-scores of IDO1 gene expression across the indicated conditions in previously published studies. Z-scores were computed separately for each study. Red boxes represent PBMC RNA-Seq data (GEO Series GSE63085), while blue boxes represent skin punch microarray data (GEO Series GSE154916). The p-values were calculated using the R package limma.

We next investigated whether the metabolic changes detected in plasma originated at the site of infection. To address this, we analyzed the skin scRNA-Seq data for the expression of metabolic enzymes involved in the pathways upregulated in plasma metabolomics data in Fig. 4a **(Table S7)**. Cells within EM lesions showed evidence of metabolic reprogramming, including upregulation of IFN-γ-inducible IDO1, which catalyzes the rate-limiting step in tryptophan metabolism, and enzymes involved in sphingolipid metabolism critical for maintaining endothelial barrier integrity **(Fig. 7d)**. While IDO1 expression showed a trending but non-significant increase in PBMCs, its highest induction was observed in skin lesions **(Fig. 7e)**. Enzymes regulating lipid biosynthesis, remodeling, and post-translational modification of proteins to maintain skin barrier integrity were also differentially expressed in EM lesions **(Table S8)**. Collectively, these findings suggest a complex interplay of metabolic and immune regulatory mechanisms in the skin, reflecting a coordinated effort to balance immune activation, tissue repair, and metabolic homeostasis in response to *B. burgdorferi* infection. These results emphasize the intricate molecular adaptations occurring in LD and highlight the central role of the skin and vascular tissue in shaping systemic immune and metabolic responses.

## Discussion

The remarkable ability of *B. burgdorferi* to evade detection by the mammalian immune system underpins its survival strategy, employing mechanisms such as antigenic variation, motility, and a highly dynamic genome enriched with plasmids^2,89,90^. This study presents a comprehensive systems-level analysis of acute LD, integrating multiomic blood measurements with public datasets from the site of infection. Our findings highlight a critical disconnect between localized tissue responses and systemic immunity in LD. Plasma profiles revealed coordinated protein and metabolite signatures— linked to endothelial barrier integrity, metabolic reprogramming, and symptom severity—predominantly traced to vascular and skin-resident cells at the site of infection. In contrast, peripheral blood mononuclear cells (PBMCs) exhibited minimal activation, with classical monocytes displaying an unconventional endothelial-protective transcriptional program, deviating from their canonical inflammatory roles^91^. Together, these findings offer a more integrated and nuanced understanding of the human immune response to LD, extending the role of endothelial dysfunction—observed in other acute conditions with long-term sequelae, such as COVID-19^92^—to LD. This reconceptualizes acute LD not only as an immune condition but also one with significant vascular involvement, emphasizing endothelial stress and systemic immune evasion as central features.

Two opposing circuits of endothelial regulation emerged in acute LD. The IFN-γ-inducible cytokine axis, involving CXCL9, CXCL10, CXCL11, and IDO-1, drove endothelial inflammation, correlated with disease severity, and promoted metabolic reprogramming through the kynurenine pathway. Conversely, the protective axis—including sphingosine-1-phosphate (S1P), beta-hydroxybutyrate (BHBA), ANGPT1, and PDGFB—stabilized the endothelium and maintained vascular homeostasis **(Fig. 3a)**. These findings build on prior studies implicating IFN-γ in *Borrelia* dissemination^22^ and endothelial damage^93^, and are consistent with independent reports of sphingolipid and beta-oxidation pathway activation in early LD^77,78^, while advancing the understanding of vascular dynamics in LD. Our data indicate that IFN-γ-inducible cytokines and IDO-1 are primarily upregulated in skin-resident cells, while IFN-γ itself is produced by a unique T cell population (T cell 1) identified in skin lesions **(Fig. 7c-d)**. Similarly, type 2 cytokines (IL-13, TSLP, and CCL28) and IL-17C associated with dermal symptoms **(Fig. 2e)** likely originate from localized immune activity at the site of infection rather than PBMCs, as peripheral immune activation was minimal in our study **(Fig. 5a-d)**. Although undetectable in our meta-analysis, these cytokines may arise from innate lymphoid cells (ILCs) or skin-resident cells, such as keratinocytes, fibroblasts, and mast cells, in response to *Borrelia* infection or tick saliva^51^. Their absence in the meta-analysis may reflect the limited availability of comprehensive skin datasets in LD studies. Future studies analyzing PBMCs and EM skin samples from the same patients will be essential for definitively linking localized and systemic immune responses.

The absence of robust PBMC activation, even in cases of disseminated disease with multiple EM lesions, emphasizes the compartmentalized nature of the immune response in LD **(Fig. 5a, 6e, and S9)**. Subtle changes in PBMCs, such as modest increases in T follicular helper (Tfh) cells and non-switched memory B cells, without effector B cell activation, suggest a skewed humoral response potentially dominated by a T-cell-independent IgM response, with limited T cell help **(Fig. 5d)**. This pattern likely reflects impaired class-switch recombination to IgG, consistent with murine studies showing disrupted lymph node architecture and altered B cell function that hinder systemic pathogen clearance^18^. Tissue-derived inhibitory signals, including PD-L1 and LILRB4 identified in our skin meta-analysis **(Fig. 7c)**, may further suppress systemic immunity^37,87,88^. PD-L1, often induced by IFN-γ, suppresses T cell activation through interactions with PD-1, while LILRB4 inhibits antigen presentation and pro-inflammatory cytokine production via its immunoreceptor tyrosine-based inhibitory motifs (ITIMs). Soluble forms of these proteins in plasma may extend their immunosuppressive effects systemically^94,95^. These inhibitory signals are reinforced by a suite of slow-resolving proteins, including PRDX1, SH2B3, and CLEC4G, which persist until T3 **(Fig. 2a-b)**, likely sustaining an inhibitory milieu. IDO-1-mediated tryptophan metabolism may further amplify this environment by producing kynurenine, which inhibits T cell proliferation, promotes regulatory T cell (Treg) differentiation, and suppresses effector T cell function^96-99^. Whether these pathways are driven by *Borrelia*, tick saliva, or host adaptation to minimize tissue damage remains unclear. Their convergence, however, suggests a coordinated mechanism that suppresses systemic immunity while facilitating pathogen persistence. This highlights the importance of investigating tissue-derived inhibitory pathways as potential targets for improving LD diagnostics, therapeutics, and enhancing *Borrelia* clearance. Although we did not directly assess antigen-specific responses in PBMCs, future studies incorporating such assays will be important for determining the extent to which pathogen-specific immunity is engaged systemically, and how this relates to the compartmentalized responses we observe.

The robust peripheral immune activation observed in one individual with 35 EM lesions and high fever sharply contrasted with the predominantly localized immune responses in most LD cases **(Fig. 6a-e)**. This individual exhibited marked elevations in inflammatory intermediate monocytes, effector T cells, and pro-inflammatory cytokines, including IFN-γ-inducible chemokines (CXCL9, CXCL10, and CXCL11) and IL-15RA, far exceeding levels seen in others. Vascular markers such as ANGPT2 and IL-8, associated with the release of WPBs by vascular endothelial cells during severe inflammation or stress, were also highly elevated, suggesting severe vascular dysfunction potentially driving systemic immune engagement^83^. The underlying reasons for such a strong peripheral response remain unclear but may involve host genetic predispositions, pathogen factors, duration of infection, bacterial load, the extent or route of dissemination, or multiple independent tick bite sites. It is plausible that the balance between inhibitory (e.g., PD-L1, LILRB4) and activating (e.g., IFN-γ, CXCL9/10/11, MIP1α) signals from skin-resident cells modulates the threshold for PBMC activation. In this individual, the intensity of local signaling likely tipped this balance, engaging the peripheral immune system. In most cases, however, this threshold remains unmet, or other mechanisms suppress PBMC activation. Similar patterns in other zoonotic skin infections^100^ suggest that skin immunity may foster peripheral immunosuppression in such contexts. Notably, ANGPT2 elevation was unique to this individual **(Fig. 2d)**, as other patients with multiple EM lesions exhibited stable ANGPT2 levels despite elevated ANGPT1, a Tie2 agonist that promotes vascular stability^41^. This indicates that vascular destabilization, while possibly facilitating pathogen dissemination, is not universally involved in extensive dissemination. Together, these findings underscore the interplay between vascular and immune pathways in modulating LD severity, with robust peripheral immune activation emerging as a hallmark of more severe disease. Although immune cell activation in this individual decreased by T2 **(Fig. 6e and S9)**, this subject dropped out of the study thereafter, precluding further assessment of immune resolution or the long-term implications of such robust peripheral activation.

Our study also highlights the diagnostic potential of circulating immune signatures. Standard antibody-based diagnostics for LD are limited by poor sensitivity during early infection, a critical window when treatment is most effective, underscoring the need for alternative approaches^13^. LASSO regression models trained on immune protein data from Olink Inflammation and Immune Response panels achieved moderate accuracy at diagnosis (AUC ∼0.68) **(Fig. S1f)**. In contrast, integrating the full proteomics and metabolomics datasets markedly enhanced accuracy (AUC = 0.94) and identified a reduced panel of 12 analytes with comparable discriminatory power (AUC = 0.92) **(Fig. 3b-d)**. These analytes reflect the interplay of inflammatory and vascular responses, offering a promising foundation for diagnostics leveraging tissue and vascular immune markers detectable in blood. Future studies must validate the specificity of these biomarkers for LD relative to other infectious and inflammatory conditions to determine whether they have clinical utility as LD-specific diagnostics or broader applicability to other conditions.

In summary, this study reveals the compartmentalized nature of immune responses in LD, with robust tissue activity and vascular involvement contrasted by minimal peripheral activation. This localized immune regulation—shaped by tissue-derived inhibitory signals such as PD-L1, LILRB4, slow-resolving proteins, and metabolic reprogramming—may limit systemic immune activation, complicating diagnosis and potentially contributing to incomplete pathogen clearance in some individuals. While antibiotic therapy effectively resolves acute symptoms in most cases, insufficient systemic immune activation may hinder full clearance and contribute to persistent symptoms in a subset of patients—a possibility that warrants further investigation. Although our study focused on acute LD, the immunologic features observed here may offer mechanistic insights into other post-infectious syndromes characterized by long-term, subjective symptoms, such as chronic fatigue syndrome^12^. Further studies are needed to determine whether similar patterns of localized immune activity and limited peripheral responses contribute to symptom persistence across these conditions. Our findings highlight the need to refocus efforts on tissue immunity and its interplay with vascular responses in LD, both as diagnostic avenues and therapeutic targets. Future vaccine strategies that elicit robust systemic and tissue-resident immunity to *Borrelia* may hold promise for improving patient outcomes. Together, these insights advance our understanding of LD pathophysiology, providing a foundation for biomarker discovery and guiding the development of more effective diagnostics and interventions to address the complexities of this disease.

## Acknowledgments

This work was supported by the Department of Defense USAMRAA, through the Tick-Borne Disease Research Program under Award No. W81XWH2110664, the Wilke Family Foundation, the Steven and Alexandra Cohen Foundation, and a fellowship from the Global Lyme Alliance to HH and CR. The authors thank Jim Heath, Rick Edmark, and Kim Murray for sharing data from the Olink Cardiovascular, Organ Damage, and Metabolic panels. We are especially grateful to the patients who contributed samples for this study, whose participation made this research possible. Opinions, interpretations, conclusions, and recommendations are those of the authors and are not necessarily endorsed by the USAMRAA.

## Author Contributions

N.S. and C.R. conceptualized the study and designed experiments. C.R. performed flow cytometry and PBMC scRNA-Seq experiments, led the analysis of flow cytometry, Olink proteomics, metabolomics data, and skin scRNA-Seq meta-analysis, prepared most figures, and drafted the initial manuscript. A.B. led the analysis of PBMC scRNA-Seq data, identified metabolic enzyme-encoding genes upregulated in the skin, contributed to community analysis in collaboration with N.R. and L.P., conducted modeling of integrated proteomics and metabolomics datasets, curated code for deposition, prepared figures, and contributed to writing. H.H. generated the Olink inflammation and immune response panel data and assisted with panel design, execution and analysis of flow cytometry. L.P. conducted community analysis. M.E.B. coordinated study logistics, established protocols, and facilitated project management. D.C. processed blood samples into plasma and PBMCs and provided clinical metadata. A.S.A. assisted with panel design, execution and analysis of flow cytometry. B.S. processed metabolomics data. K.S. and K.W. contributed to Olink panel experiments. P.T. prepared PBMC scRNA-Seq libraries. C.L. preprocessed PBMC scRNA-Seq data. G.P.W. recruited, clinically assessed patients, contributed to the interpretation of clinical data, and edited the final manuscript. N.R. supervised the community analysis. L.H. initiated and led the broader systems biology of Lyme disease project, secured funding, and edited the final manuscript. N.S. supervised the study, directed research strategies, secured funding, and wrote and edited the final manuscript with input from all authors.

## Inclusion and Ethics Statement

This study was approved by the WIRB Institutional Review Board (IRB approval #10999), and written informed consent was obtained from all participants prior to enrollment. There were no gender, racial, or ethnic exclusion criteria. The demographic composition of the study population reflected that of the general population in the New York metropolitan area. PBMC samples have been largely depleted following flow cytometry and scRNA-Seq experiments, but limited volumes of plasma remain from a subset of patients and may be shared upon reasonable request and institutional approval.

## Data availability

scRNA-Seq data generated in this study will be deposited in the Gene Expression Omnibus (GEO) database, and proteomic data will be made available in the PRIDE repository before or immediately upon publication. Metabolomics data will be deposited by Metabolon in a publicly accessible repository, with details provided in the final manuscript. All raw data and associated metadata are available upon request from the corresponding author.

## Code availability

Custom code developed for this study is available at https://github.com/subramanian-group/Lyme. The repository is currently private but will be made publicly accessible upon publication and made available to reviewers during peer review. A permanent DOI will be provided in the final manuscript.

## Methods

### Study Enrollment

Adults 18 years of age or older were recruited into this study from the Lyme Disease Diagnostic Center located in Westchester County, New York State. The demographic composition of the study population reflected that of the general population in the New York metropolitan area. There were no gender, racial, or ethnic exclusion criteria. Pregnant or immunocompromised individuals were excluded. A diagnosis of Lyme disease (LD) was made by visual inspection of the skin for an erythema migrans (EM) lesion by study investigators, along with other objective clinical manifestations and serologic confirmation, which included a positive C6 ELISA and a two-tiered Western blot test, and in some cases a positive blood culture for *Borrelia*. EM was defined in accordance with CDC criteria. Preferred study participants were those who had not yet received antibiotic treatment for LD at the time of enrollment. Patients with a prior history of LD within the past 12 months or with continuing symptoms were excluded.

Additional exclusion criteria for both patients and controls included a history of fibromyalgia, chronic fatigue syndrome, or traumatic brain injury prior to LD onset, a prolonged history of undiagnosed or unexplained somatic complaints (e.g., fatigue or musculoskeletal pain), or medical or psychiatric conditions that could interfere with outcome evaluation or follow-up. These included morbid obesity (BMI ≥45), sleep apnea and narcolepsy, autoimmune disease, uncontrolled cardiopulmonary or endocrine disorders, recent malignancy, liver disease, major psychiatric illness, or substance abuse. Healthy controls were also excluded if they had a prior history of LD or were seropositive by two-tier serologic testing at the baseline visit. Control subjects were recruited during the same LD season as patients with EM to minimize temporal variation in exposure and immune profiles.

Symptom questionnaires were collected from study participants at the initial diagnosis (T1), 6 month (T3), and 1 year (T4) visits. Subjects rated the severity of 12 symptoms known to be associated with LD using a visual analogue scale. This was followed by a physician interview to determine the relevance of those symptoms to LD or to an alternative diagnosis. All study procedures were reviewed and approved by the WIRB Institutional Review Board (IRB approval #10999).

### Blood Processing

Plasma was isolated from fresh whole blood by centrifugation, aliquoted and stored at -80°C. PBMCs were isolated using Ficoll density gradient centrifugation and viability and cell count were recorded.

PBMCs were cryopreserved using a freezing medium containing 10% DMSO in FBS and stored in liquid nitrogen. Samples were shipped on dry ice from NYMC to the Institute for Systems Biology (ISB). Upon receipt plasma was again stored at -80°C, and PBMCs were stored in liquid nitrogen until use.

### Plasma Proteomics

#### Plasma protein measurements by Olink Proteomics proximity extension assay (PEA)

The abundance of 460 unique proteins in the plasma was analyzed using five Olink Multiplex Assay Panels—Inflammation, Immune response, Cardiovascular, Metabolism, and Organ Damage—and quantified by real-time PCR using the Fluidigm BioMark™ HD platform according to the manufacturer’s instructions. The proximity extension assay (PEA) method relies on two oligonucleotide-labeled antibody probes that bind to each target protein. When two antibody probes are in close proximity, a proximity-dependent DNA polymerization event generates a unique PCR target sequence, which is subsequently detected and quantified by real-time PCR. Briefly, plasma samples were thawed and centrifuged to remove aggregates. Incubation reagents for oligo-conjugated antibody binding were thawed, vortexed and briefly centrifuged. Assay mastermix was transferred to a 96-well plate using reverse pipetting, and 1uL of plasma was added to the mastermix-containing plate using a multichannel pipette. The plate was sealed and incubated overnight at 4°C (17 hours). The extension mix was then added to each well using reverse pipetting, followed by gentle mixing and centrifugation, and the PEA extension reaction was completed in a thermocycler. Detection reagents were thawed and mixed, and 7.2uL of detection mix was added to each well of a new 96-well detection plate. Subsequently, 2.8 uL of sample from the extension plate was transferred to the detection plate. For real-time PCR, 5 uL of the sample mixture was loaded into the sample inlet wells of a primed microfluidic real-time PCR chip (Biomark 96.96 Dynamic Array, Fluidigm). Similarly, 5 µL of primer mix was added to the corresponding wells of the array, following the Fluidigm protocol. Analytes were detected by running the chip on the BioMark™ HD system with the following thermal program: Thermal mix (50°C for 2 min; 70°C for 30 min; 25°C for 10 min), Hot-start (95°C for 5 min), PCR Cycle for 40 cycles (95°C for 15 s; 60°C for 1 min).

#### Olink Data Processing

Olink data were normalized and converted to Normalized Protein eXpression (NPX) values by following the manufacturer’s protocol (www.olink.com). The resultant NPX values were batch normalized using an empirical Bayes framework implemented in the ComBat function of the sva R package by defining each run as a unique batch and modeling age, gender, and infection condition as covariates^101^.

#### Olink Data Analysis

Differentially expressed proteins were computed by fitting linear models using the R limma package^102^. Models were fit including age, gender, and seropositivity as additional response variables to account for their effect on protein levels. P-values were adjusted using the Benjamini-Hochberg correction^103^, and a significance threshold of p.adj <= 0.05 was set to define differentially expressed proteins. We clustered pairwise differences between patient timepoints by ward.D2 linkage. The resulting hierarchical tree was cut at k=2 to reveal the fast- and slow-resolving clusters. Gene set enrichment analysis was conducted using the clusterProfiler R package^104^ using MsigDB Hallmark, C2, C4, and C7 datasets^105^. Correlations and their significance were determined using the rcorr R package.

### Predictive modeling of plasma proteomics data

LASSO regression models were built to classify patient samples using two different approaches. The first set of models was fit to NPX values from the Olink Inflammation and Immune Response panels using the glmnet R package^106^. Monte Carlo nested cross-validation was used to assess model performance with 500 iterations. In each iteration, 80% of the data was randomly selected as the training set, with hyperparameter tuning (i.e., selection of the LASSO penalty) performed using 5-fold internal cross-validation on the training set. The remaining 20% of the data was used as the test set to calculate the Area Under the Curve (AUC) for each iteration. The mean AUC across all test set iterations was reported, along with the corresponding mean AUCs for the models built using the full dataset and the training set.

The second set of models was fit to combined Olink and Metabolon data using the glmmLasso R package^107^ to account for within-subject correlations between paired samples. A random intercept was included for each subject to control for baseline differences. Monte Carlo cross-validation with 500 iterations was used to evaluate model performance. In each iteration, the dataset was randomly split into an 80% training set and a 20% test set, ensuring that both time points (T1 and T3) for each patient were retained within the same partition. The LASSO penalty was applied to the fixed effects, while the random intercept controlled for subject-level variability.

For further analysis, a reduced model was constructed using the 77 analytes identified in the community analysis. To refine the model, the analytes were further reduced to 12 key predictors that appeared in at least 300 of the 500 simulations conducted during Monte Carlo cross-validation. Model performance was assessed by calculating the AUC for each iteration using the test set predictions. The median ROC curve across all iterations was extracted, along with the 95% confidence interval (CI) for the True Positive Rate (TPR) at each False Positive Rate (FPR) threshold.

### Metabolomics

#### Measurement of circulating metabolites using Metabolon

Plasma samples were transferred to Metabolon (Morrisville, NC, USA) for metabolomic assays. Samples were processed on Metabolon’s Global Metabolomics platform using high-performance liquid chromatography (HPLC)-mass spectrometry (MS) platform. For most samples, 250 ul of plasma in EDTA was aliquoted and shipped on dry ice to Metabolon. Six samples had less plasma available and smaller amounts were shipped (100-225 ul). Metabolon performed QA/QC in their CAP/CLIA-certified laboratory before analyzing samples as previously described^108^. Briefly, samples were run on four independent instruments that consist of a Waters ACQUITY ultra-performance liquid chromatography (UPLC) and Thermo Scientific Q-Exactive high resolution/accuracy mass spectrometer interfaced with a heated electrospray ionization (HESI-II) source and an Orbitrap mass analyzer operated at 35,000 mass resolution. Quality control samples were included in each batch and were consistent across batches. Metabolon completed the initial data quality analysis, quantification, and compound identification.

#### Metabolon Analysis

Metabolites with fewer than six observations across the entire dataset were removed from downstream analysis. Data were then normalized for downstream analysis by dividing each metabolite by its median and log scaling the resulting values. Log fold changes (LFC) of metabolites across time points were computed as LFC = mean(log(A)-log(B)), where A and B are paired vectors of patient metabolite at the time points being compared. For each metabolite, full and reduced linear models shown below were compared using a likelihood ratio test to test for the effect of time pairwise while addressing covariates. These models were fit in R using the lm function, and the likelihood ratio test was computed using the lrtest function from the lmtest R package^109^.

Full Model: log(Metabolite Value) ∼ Time + Sex + Patient ID + Age + BMI + Pulse

Reduced Model: log(Metabolite Value) ∼ Sex + Patient ID + Age + BMI + Pulse

Significant metabolites from the likelihood ratio test were used as input for pathway enrichment analysis using the R package FELLA^110^.

### Flow cytometry-based immunophenotyping

#### Sample Processing

Cryopreserved PBMCs were rapidly thawed and stained using the antibodies indicated in **Table S1**. Briefly, cells were first stained with a viability stain. Fc receptors were blocked using human IgG and staining for surface markers was performed at 4°C for 60 minutes protected from light. Cells were fixed and permeabilized, and intracellular markers were stained using the BD FoxP3 buffer kit (cat# 560098) according to the manufacturer’s instructions. Cells were then stored overnight in BD Cytofix buffer (cat# 554722) at 4°C protected from light. Prior to acquisition, cells were diluted 1:1 in FACS buffer (1X PBS supplemented with 1% BSA, 2 mM EDTA, and 0.1% sodium azide), and flow cytometry was performed at the University of Washington Flow Cytometry core facility using the BD LSRII.

#### Data Processing

Flow Cytometry data were processed as shown in **Fig. S4**. Data were imported into R using the flowCore package^111^. Artifacts were removed using the nmRemove function from the flowDensity package^112^, and samples with less than 30k cells were removed. FSC-low SSC-low debris from every sample was removed by modeling FSC and SSC as a mixture of 3 Gaussians using the GMM function from the ClusterR R package^113^ and defining the debris cutoff as the point where the lowest FSC Gaussians intersected with the middle Gaussian times a scaling factor. Doublets were then removed by fitting a robust linear model to the FSC-A vs. FSC-H data and removing events whose residuals were greater than 3.5 median absolute deviations. Samples were compensated for spectral overlap using FlowJo software. Single color beads defined positive peaks for each marker. The spectral compensation function was employed to utilize signals from detectors not intended for primary data acquisition. Spillover into these detectors was used to separate individual colors more precisely and further improve spectral unmixing (www.flowjo.com). The resultant spectral overlap matrix was then manually adjusted to correct for overcompensation and differences between beads and cells. Fluorescent channels were asinh transformed using cofactors determined by the flowVS R package^114^. Next, the flowClean R package^115^ was used to remove time periods with abnormal fluorescence due to stream instability. Dead cells were removed by defining a dead cell cutoff for each plate as the median value of viability gates determined using the deGate function from the flowDensity R package^112^. The median was chosen to avoid issues with batch application of the deGate function on individual samples. Finally, samples were manually gated for top populations (e.g. CD3+ for T cells) using gates defined by the deGate^112^ and ddtw^116^ algorithms as guides to help determine gate placement for each sample.

#### Clustering and Gating

For the B cell, monocyte, and dendritic and natural killer cell panels, populations were determined using the CATALYST pipeline^117^. Cells were clustered using self-organizing maps implemented in the flowSOM package^118^. The resulting clusters were merged using ConsensusClusterPlus^119^, and these final clusters were manually merged and annotated to resolve the immune cell clusters shown in **Fig. S5**. The clusters derived from the T cell panel did not segregate cleanly into known populations, likely because of the combinatorial nature of T cell lineage, memory, and effector function markers. To resolve populations in the T cell panel, gates were automatically determined for each marker using the flowDensity R package^112^. Using these gates populations were manually annotated to reveal the T cell populations shown in **Fig. S5**.

#### Downstream Analysis

Differential abundance testing was conducted using the diffcyt package with the testDA_edgeR option^120^. Correlations were computed using the rcorr function from the Hmisc package [Hmisc 2021]. PCA was computed on the z-scores of cell type proportions of total cells. Some measurements were missing from these data and their values were imputed using the ‘mice’ package^121^ prior to computing the PCA.

### Transcriptomics

#### PBMC scRNA-Seq

##### Data acquisition

Sixteen PBMC samples were processed using eight 10x Genomics Chromium bead libraries by sample pooling. For each library, two genetically distinct samples were pooled at equal cell concentrations, targeting 4000 cells per sample. These libraries were sequenced on the NovaSeq platform (Illumina) by the Northwest Genomic Center at University of Washington. The read configuration was 28 cycles for Read 1, 10 cycles for the i7 index read (Index 1), 10 cycles for the i5 index read (Index 2), and 90 cycles for Read 2.

##### Sequence alignment

FASTQ file reads were aligned, cells identified, and genes counted using CellRanger (v3.0, 10x Genomics) with GRCh38 as the reference genome. The expected number of cells per sample was set to 8000. Pooled samples were demultiplexed using Vireo (v0.5.8)^122^ with a list of 7.4M common polymorphisms (minor allele frequencies > 0.05) derived from the 1000 Genomes Project (https://sourceforge.net/projects/cellsnp/files/SNPlist/). From each library, we obtained 6182 ± 966 cells, 139000 ± 16000 reads per cell, 1802 ± 110 genes per cell, and a sequencing saturation of 88.5 ± 1.4 percent.

##### Data analysis

Low-quality cells were removed using the quickPerCellQC command from the R package Scuttle ^123^. This identifies and removes low-quality cells based on their library size and the number of detected features. In addition, cells with a high proportion of mitochondrial transcripts were also removed, resulting in approximately 188,000 cells. The gene-by-cell matrices of the individual samples were normalized and then integrated using canonical correlation analysis (CCA) in Seurat ^124,125^. After integration, cells with fewer than 500 unique features were removed, resulting in 42,000 cells. The integrated data were subjected to dimension reduction using PCA and UMAP. The first 30 PCA dimensions were utilized to detect shared nearest neighbors (SNN)^126^. The neighborhoods were resolved into separate clusters using the Louvain algorithm implemented in Seurat^127^. The identities of the clusters were determined by mapping them onto the Monaco bulk-RNA sequencing dataset using the SingleR package^128,129^. Within each cluster, cells from the same patient were pseudo-bulked to account for within-patient correlation before performing the differential expression analysis ^130^. The differentially expressed genes were identified by comparing T1 samples to T4 samples (reference) in a pairwise manner using DESeq2 ^131^.

#### Analysis of public bulk gene expression data

Public data for GSE154916^20^ and GSE84479^132^ were downloaded from the Gene Expression Omnibus (GEO) using the GEOquery R package^133^. Count data for GSE63085 was obtained from the GREIN repository^134^, and metadata was obtained from GEO **(Table 2)**. Differential expression was computed using the limma R package^102^. RNA-Seq counts in GSE63085 were filtered by removing genes with a max count of less than 2, and then normalized using voom prior to differential expression with limma^135^. Cell type enrichment analysis from gene expression data **(Fig. S10a)** was done using the R package xCell^85^. Cohen’s d was calculated as the standardized measure of differences in cell-type enrichment scores between experimental groups (shown along the lower x-axis in Fig. S10a). For each cell type, a linear model was fitted with enrichment scores as the outcome, and the corresponding pairwise contrasts were estimated using the emmeans package. Cohen’s d was then computed by dividing each contrast estimate by the pooled residual standard deviation from the model.

#### Analysis of public skin scRNA-Seq data

##### Preprocessing, integration, and cell subset annotation

scRNA-Seq data from GSE169440^86^ were analyzed using Seurat v4^125^. Each sample was independently normalized and the 2000 top variable features were determined for each dataset. The features consistently variable across samples were then selected for integration of the datasets. Integration was performed according to Stuart *et al.* 2019^124^. The resulting integrated data were log scaled and used for PCA and UMAP dimension reductions. The first 30 PCA dimensions were used to identify shared nearest neighbors (SNN)^126^, and the SNN graph was clustered using the modularity optimization-based clustering algorithm implemented in Seurat. Marker genes for each cluster were determined using the original unintegrated counts, and cluster identities were manually annotated. A summary of marker genes and annotations is shown in **Fig. S11**. Each of the clusters was then subsetted and the same clustering approach was implemented on each cluster individually. The resulting sub-clusters and their marker genes are shown in **Fig. S12 and S13**.

##### Differential cell abundance analysis in skin samples

The clusters of skin cells present in the scRNA-Seq data of EM lesions, GSE169440^86^, were manually annotated as described above. For each cell cluster, a generalized linear model (GLM) was fitted to assess differential abundance, after including terms for subject ID and skin condition (normal or EM). The p-values were adjusted using the false discovery rate (FDR) method.

##### Differential gene expression analysis in skin samples

Within each cluster of skin cells, cells from the same patient were pseudo-bulked to account for within-patient correlation before performing the differential expression analysis^130^. The differentially expressed genes were identified by comparing EM samples to normal skin samples (reference) in a pairwise manner using DESeq2^131^.

##### Selection of differentially expressed metabolic enzyme genes

We focused on genes encoding enzymes in the arginine biosynthesis, sphingolipid metabolism, and energy metabolism pathways on KEGG. Initially, cross-species orthologs were included among the hits. Subsequently, the KEGGREST package (Tenenbaum and Bioconductor package maintainer 2024) was utilized to match the orthologs to a singular entry in our list of differentially expressed genes. Only genes with an adjusted p-value below 0.05 were kept for further analysis.

### Community analysis

#### Community detection and interomic correlation analysis

Community analysis was performed as previously described^136^. Briefly, we created correlation networks at time T1 and T3. Prior to analysis, proteomic and metabolomic datasets were filtered for analytes with less than 50% missingness; metabolites were additionally filtered to exclude unannotated metabolites. Spearman’s ρ values were calculated between proteomic and metabolomic datasets, followed by p-value adjustment for multiple hypothesis testing using the method of Benjamini and Hochberg. We chose an adjusted p-value of 0.05 as our significance level cutoff. Community analysis was performed using the Girvan and Newman method (2002), with the final network having the highest modularity after edge cutting. Only interomic, and not intraomic, correlations were used for the community analysis. Network analysis and community detection were performed using the Igraph R package^137^. Community eigenvectors were derived using the first principal component for all analytes in the identified communities. Missing data were imputed using KNN, centered, and scaled prior to generating eigenvalues. All analyses were performed using R (v. 4.3).

#### Differential expression of community metabolites

Metabolon data that were normalized earlier for the likelihood ratio test were filtered for antibiotic-naive patients and analyzed to identify community metabolites that were differentially expressed between time points T1 and T3. Metabolites with fewer than five paired observations were removed from downstream analysis. A linear mixed-effects model was fit for each metabolite, using time as a fixed effect and patient as a random effect to account for repeated measures. The resulting p-values were adjusted for multiple testing using the Benjamini-Hochberg method. Metabolites with an adjusted p-value (FDR) ≤ 0.05 were considered significantly differentially expressed.

#### Differential expression of community proteins

Olink data that were normalized and batch-corrected earlier for the limma analysis were filtered for antibiotic-naive patients and analyzed to identify community proteins that were differentially expressed between time points T1 and T3. For each protein in the five Olink panels a linear mixed-effects model was used to assess the effect of time on Normalized Protein Expression (NPX) values, with time as a fixed effect and patient as a random effect. The resulting p-values were adjusted for multiple testing using the Benjamini-Hochberg method. Proteins with an adjusted p-value (FDR) ≤ 0.05 were considered significantly differentially expressed.

#### Correlation of community factor loadings with symptom score

Loadings indicating the contribution of each factor (metabolite or protein) to the respective principal component were calculated. To assess the clinical relevance of the factors driving the separation in the principal component space, we examined the correlations between average symptom scores for each patient and the factor loadings. Spearman’s rank correlation was employed to capture potential nonlinear relationships, and p-values were adjusted using the Benjamini-Hochberg method to account for multiple comparisons.

## Supplementary tables

**Table S1: Markers used for immunophenotyping PBMCs.** Antibodies employed for flow cytometry staining of T cells, B cells, monocytes, DCs, and NK cells are shown.

**Table S2: Annotation of immune cell populations analyzed by flow cytometry.**

**Table S3: Genes differentially expressed in patient PBMCs in scRNA-Seq.** We compared the gene expression in PBMCs at two time points within patients: T1 (acute infection phase) and T4 (one year later; reference). The table lists the differentially expressed genes by cell type (FDR < 0.05).

**Table S4: Plasma protein z-scores for all differentially expressed proteins from the Olink Inflammation and Immune Response panels at diagnosis.** Z-scores were computed on all patient and control samples at all time points. Mean_Z is the mean z-score of the patients at T1 relative to the rest of the samples. SD_Z is the standard deviation of the z-scores of the patients at T1 relative to the other samples. Outlier_Z is the z_score of the outlier patient (201455 T1) with 35 EM.

**Table S5: Plasma protein z-scores for all differentially expressed proteins from the Olink Metabolism, Cardiovascular, and Organ damage panels at diagnosis.** Column descriptions are identical to those in Table S4.

**Table S6: Metabolite z-scores for all differentially expressed metabolites in the plasma at diagnosis.** Column descriptions are identical to those in Table S4.

**Table S7: Metabolic enzyme-encoding genes differentially expressed in skin scRNA-Seq.** The table lists the enzyme-encoding genes that were differentially expressed between EM and unaffected skin in GSE169440. For each gene, the cell type differentially expressing it and the metabolic reaction it catalyzes are also shown.

**Table S8: Functions of metabolic enzymes shown in Fig. 7d.**

## Supplementary figures

**Figure S1:**
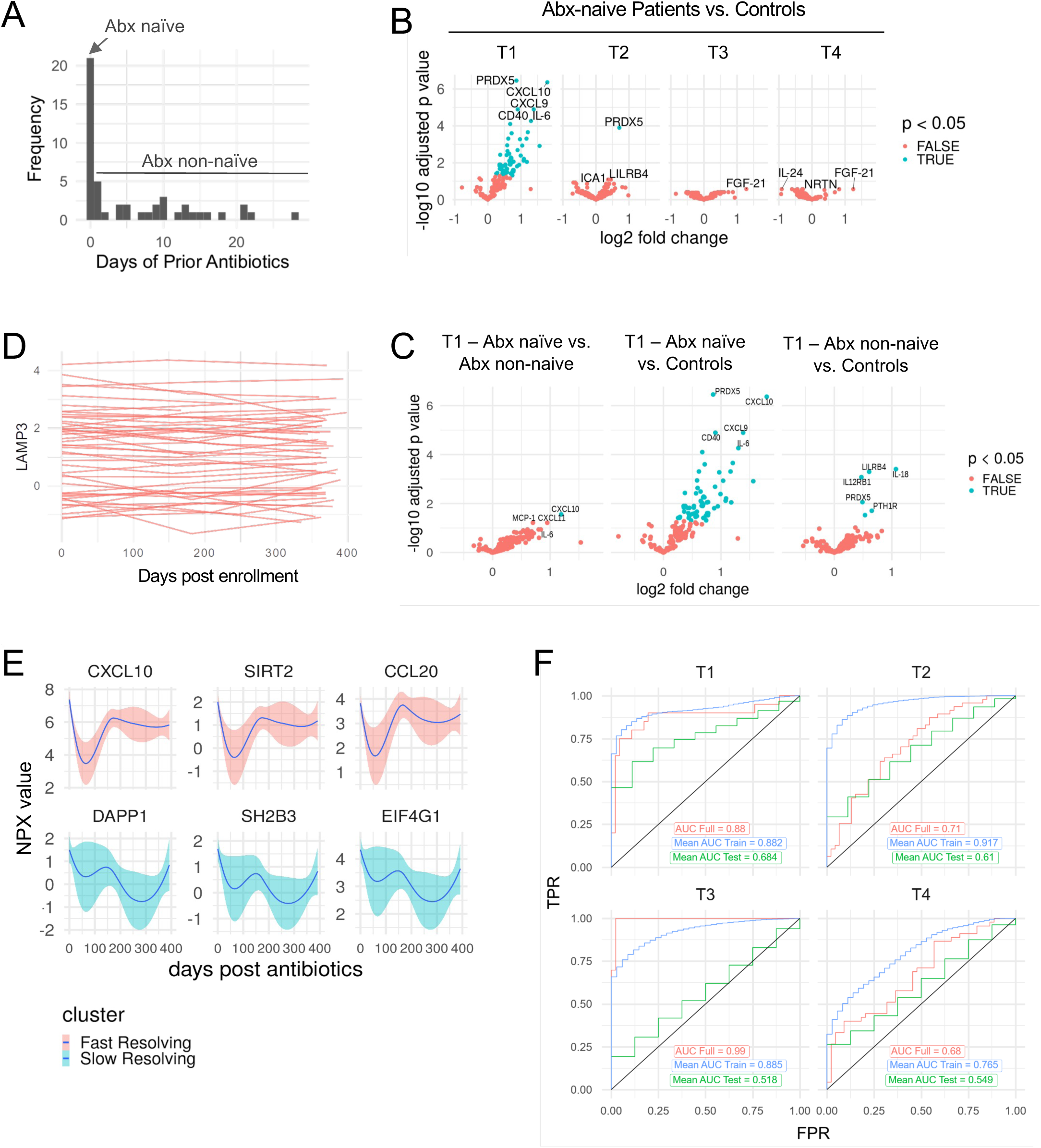
Circulating immune protein trajectories in acute disease. **(A)** Histogram of antibiotic exposure at T1. **(B)** Differentially expressed proteins between patients and controls at the indicated time points. Significant proteins (p<0.05) are highlighted in blue. **(C)** Differentially expressed proteins between the indicated groups at T1 showing significant (p<0.05) proteins in blue. Differential expression was computed using the R package limma. **(D)** Spaghetti plot of a representative Olink protein LAMP3 showing a different baseline in each healthy control. Each line represents an individual. **(E)** Line plots showing Lowess smoothed normalized protein expression (NPX) values in patients for three representative proteins each from the fast- and slow-resolving clusters. **(F)** ROC curves from a LASSO regression model with Monte Carlo nested cross-validation, classifying patients from controls at various time points using proteins from the Olink Inflammation and Immune Response panels. Test accuracy (green curve) represents the average of 500 ROC curves generated by fitting the model on randomly selected 80% subsets of the data during each iteration. Hyperparameters were optimized by internal cross-validation, and the model was implemented using the glmnet R package. Also shown are the ROC curve of the full model (red) and the average training ROC curve (blue). Time points: T1 (diagnosis), T2 (2–3 weeks), T3 (6 months), T4 (1 year).

**Figure S2:**
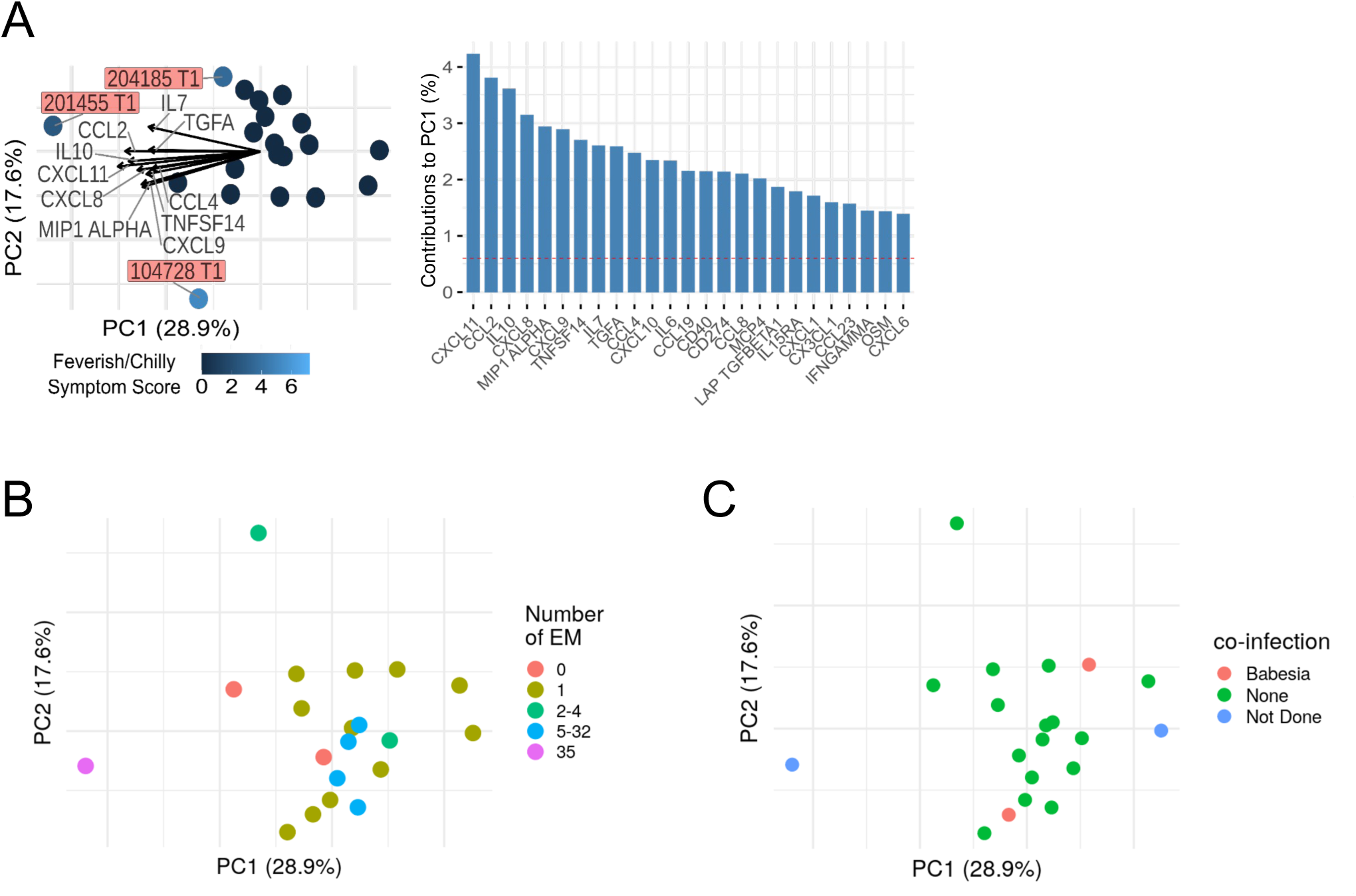
Coinfection status and number of EM do not drive changes in circulating immune proteins. **(A)** PCA of circulating proteins in antibiotic-naive patients at diagnosis colored by patient-reported severity of fever. Protein values were z-score normalized to all patients and controls in the study prior to computing the PCA. Arrows represent the top 15 factor loadings. The bar graph on the right indicates the contribution of the top loadings to PC1. **(B-C)** PCA of Olink proteomics data from antibiotic-naive patients colored by number of EM (B) and co-infection status (C). PCA was computed as in A.

**Figure S3:**
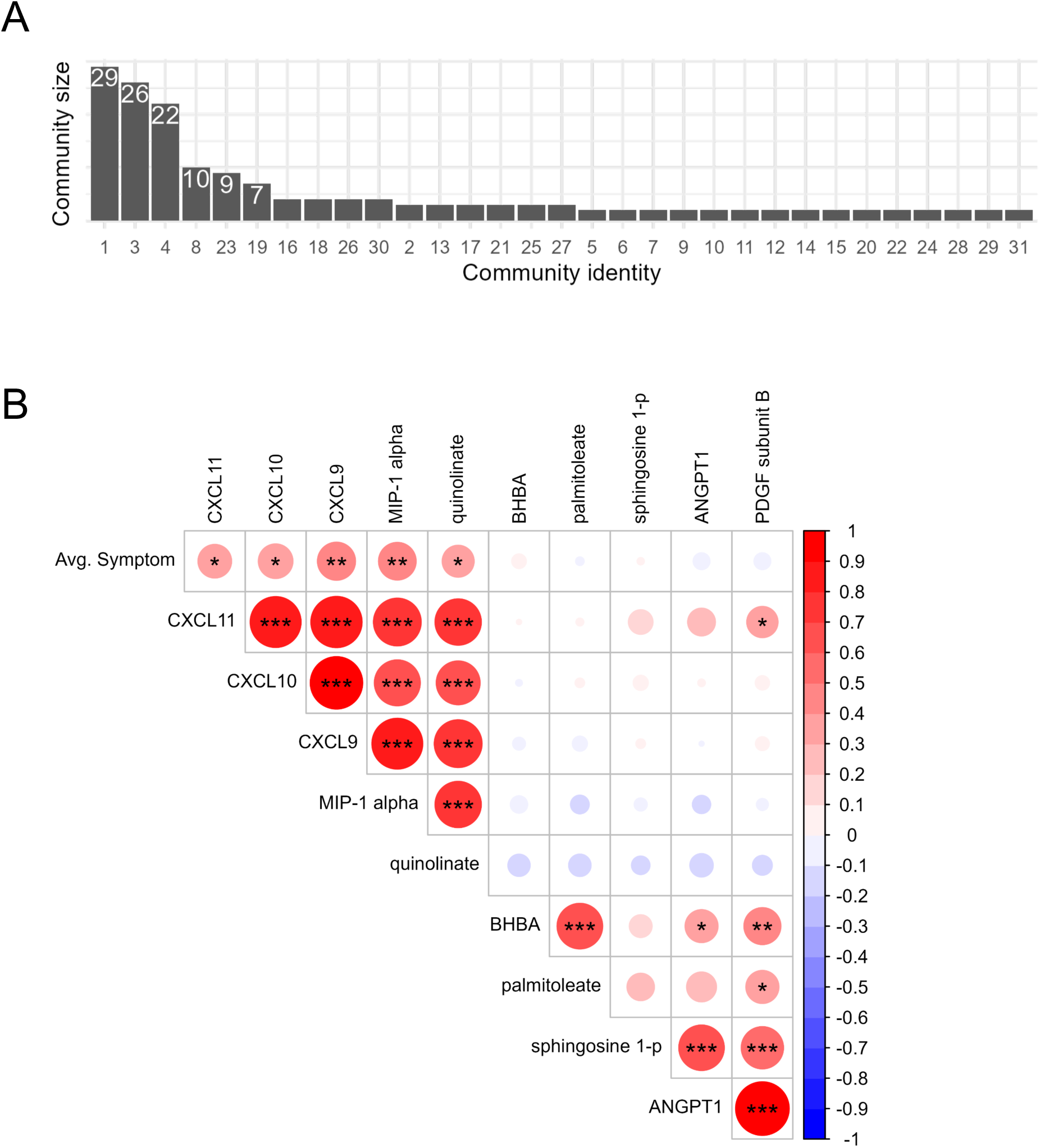
Community composition and correlation of proteins and metabolites with symptoms. **(A)** Bar plot showing the number of analytes in each community. The community identity is displayed along the x-axis, while the exact sizes are labeled on top of the bars. **(B)** Correlation matrix showing Spearman correlations among the indicated circulating proteins, metabolites, and symptom severity. Proteins and metabolites were drawn from the largest inter-omic community identified in panel A. Circle size and color denote correlation strength and direction; asterisks indicate significance (*p < 0.05, **p < 0.01, ***p < 0.001).

**Figure S4:**
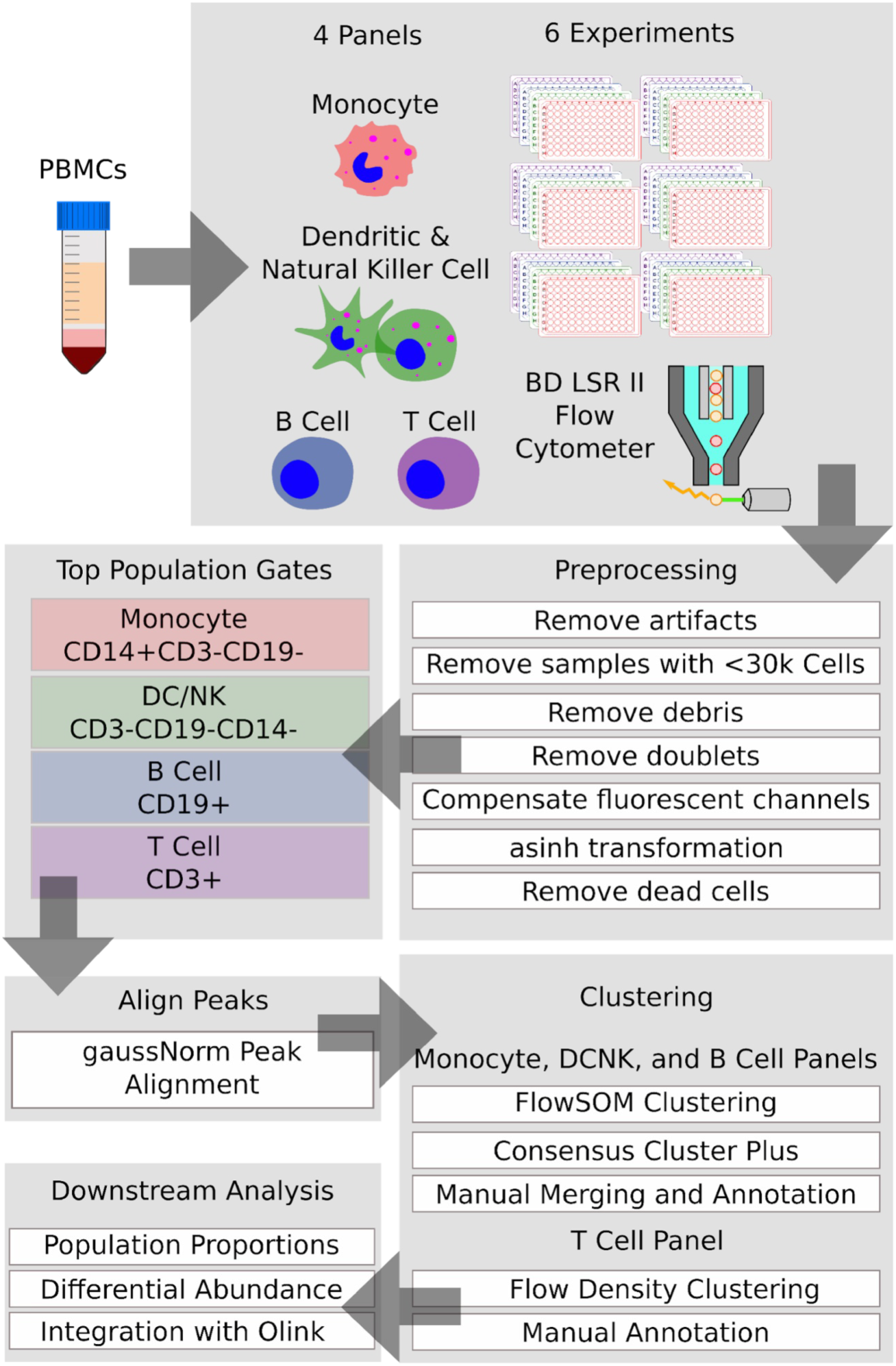
Pipeline for analysis of flow cytometry data. The flowchart details a stepwise systematic analysis of flow cytometry data including data preprocessing, unbiased clustering, and downstream analysis.

**Figure S5:**
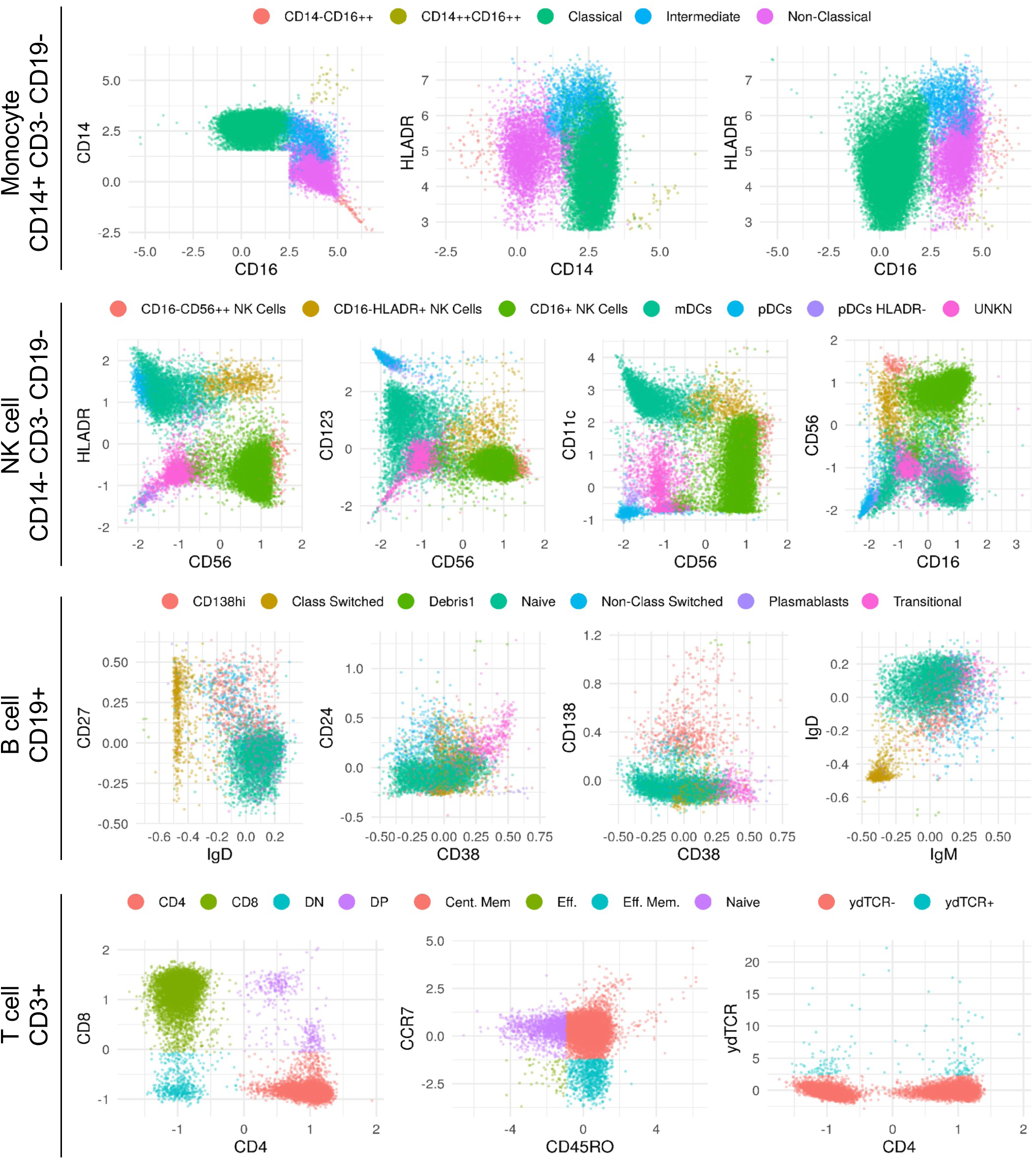
Representative plots showing clustered populations from flow cytometry data after following the analysis pipeline. Major populations of monocytes, NK, B, and T cells are shown. Activation markers and cytokines are not shown. Cent. Mem: central memory; Eff: effector; yd: gamma-delta T-cells.

**Figure S6:**
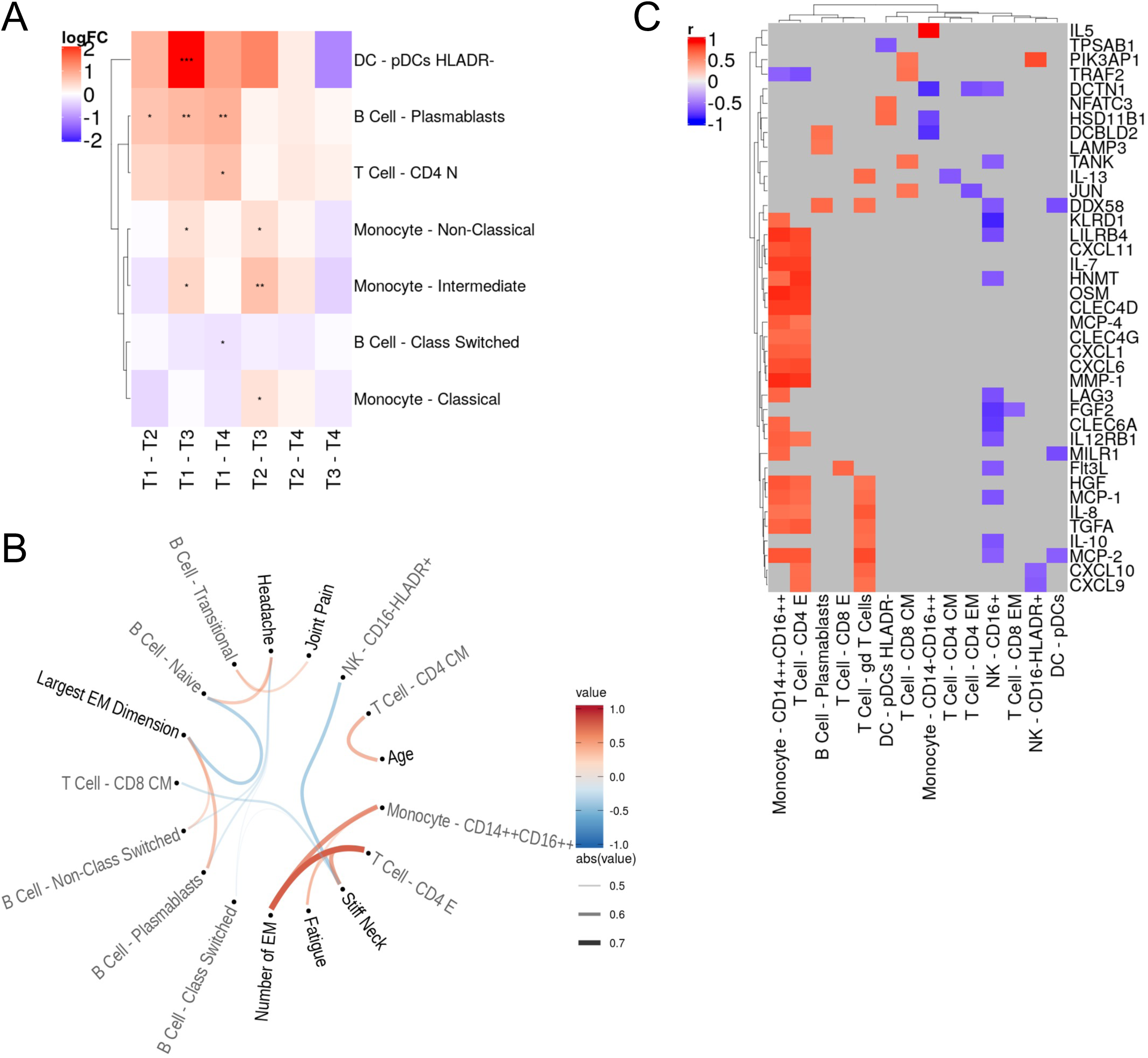
Changes in PBMC composition and their relationships with symptoms and circulating proteins in acute disease. **(A)** Heatmap of cell types with significant abundance changes over time. The diffcyt wrapper around edgeR was used to perform pairwise comparisons at the indicated time points. Only populations with significant (p<0.05) differential abundance in at least one comparison are shown. **(B)** Hierarchical edge bundling plot of significant Pearson correlations (p<0.05) between immune cell abundance and patient-reported symptom scores in antibiotic-naive patients at diagnosis. Features with similar correlation patterns are grouped together by hierarchical clustering. Line width and color indicate the strength and direction of correlation respectively. Correlations were computed using the rcorr function from the sva R package. **(C)** Heatmap of significant Pearson correlations (p<0.05) between immune cell abundance (columns) and circulating plasma proteins (rows) in antibiotic-naive patients at diagnosis. Features with similar correlation patterns are grouped together by hierarchical clustering. Correlations were computed using the rcorr function from the sva R package.

**Figure S7:**
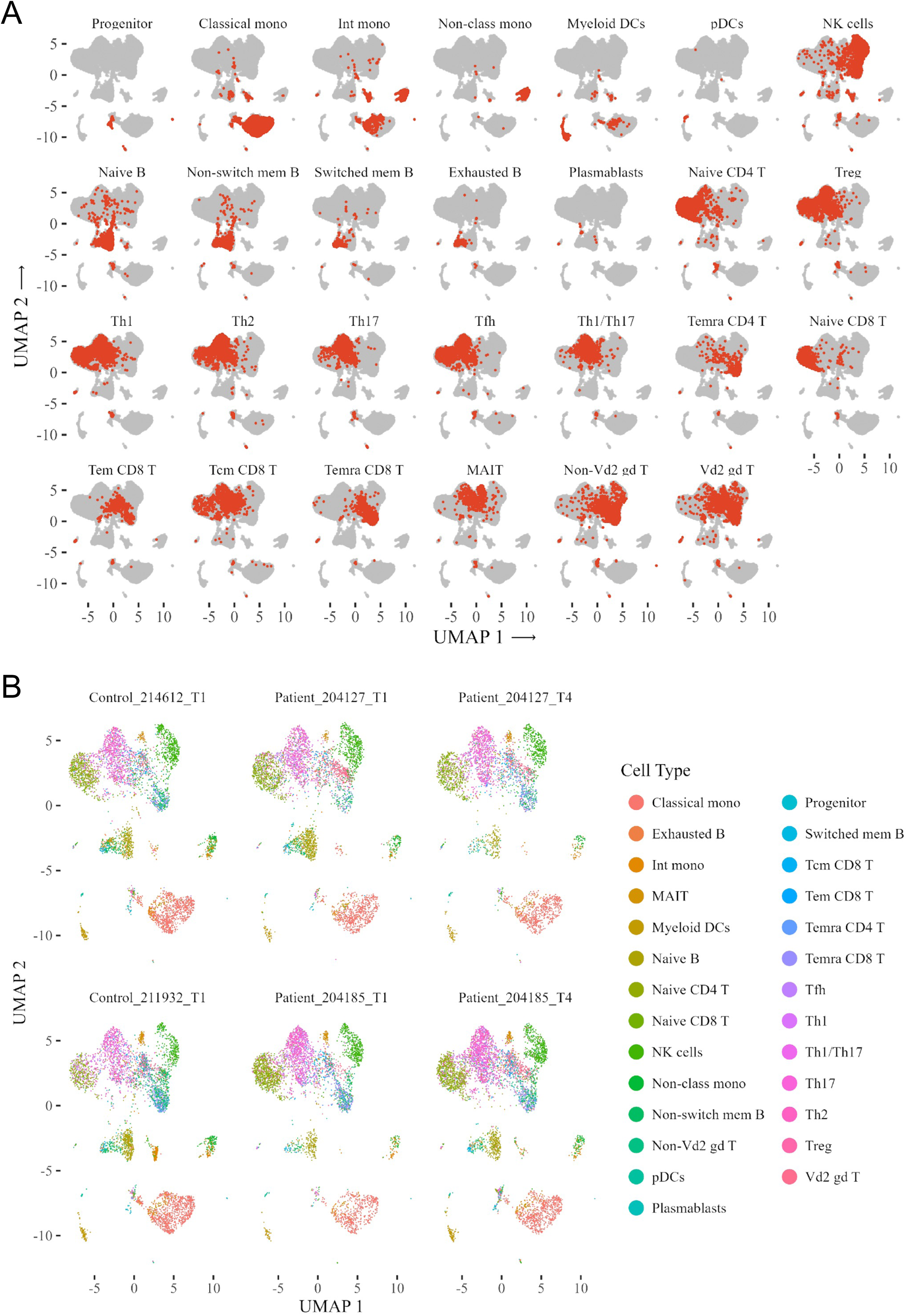
UMAPs of PBMCs after scRNA-Seq. **(A)** Small multiples of major peripheral cell populations clustered based on their scRNA-Seq UMAP distances. **(B)** UMAPs of individual PBMC samples from scRNA-Seq. UMAPs are shown for two patients with matched T1 and T4 samples, and two representative healthy controls (selected from a total of four). Data are shown to illustrate sample-level variation and overall consistency in PBMC composition across conditions. All 27 annotated cell types are displayed. Int mono, intermediate monocytes; Non-class mono, non-classical monocytes; Tfh, follicular helper T cells; Temra CD4/ CD8, terminal effector CD4⁺/CD8⁺ T cells; Tem CD8, effector memory CD8⁺T cells; Tcm, central memory CD8⁺ T cells; gdT, gamma delta T cells.

**Figure S8:**
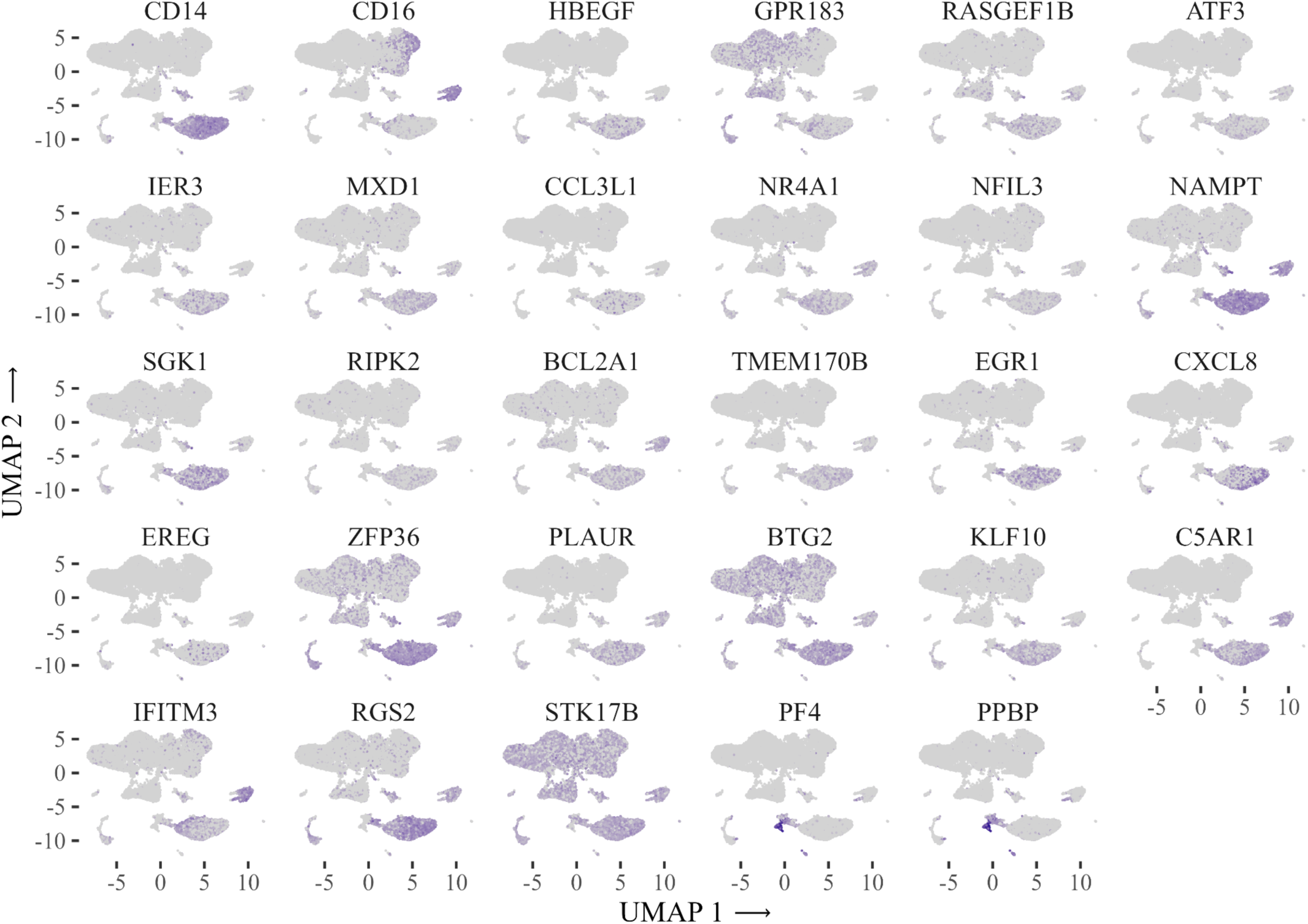
Localization of genes differentially expressed in classical monocytes. UMAP plots showing the expression of selected genes identified as differentially expressed in classical monocytes at T1 versus T4, projected across all PBMC clusters. Each panel displays scRNA-Seq expression of the indicated gene, with expression intensity shown in purple.

**Figure S9:**
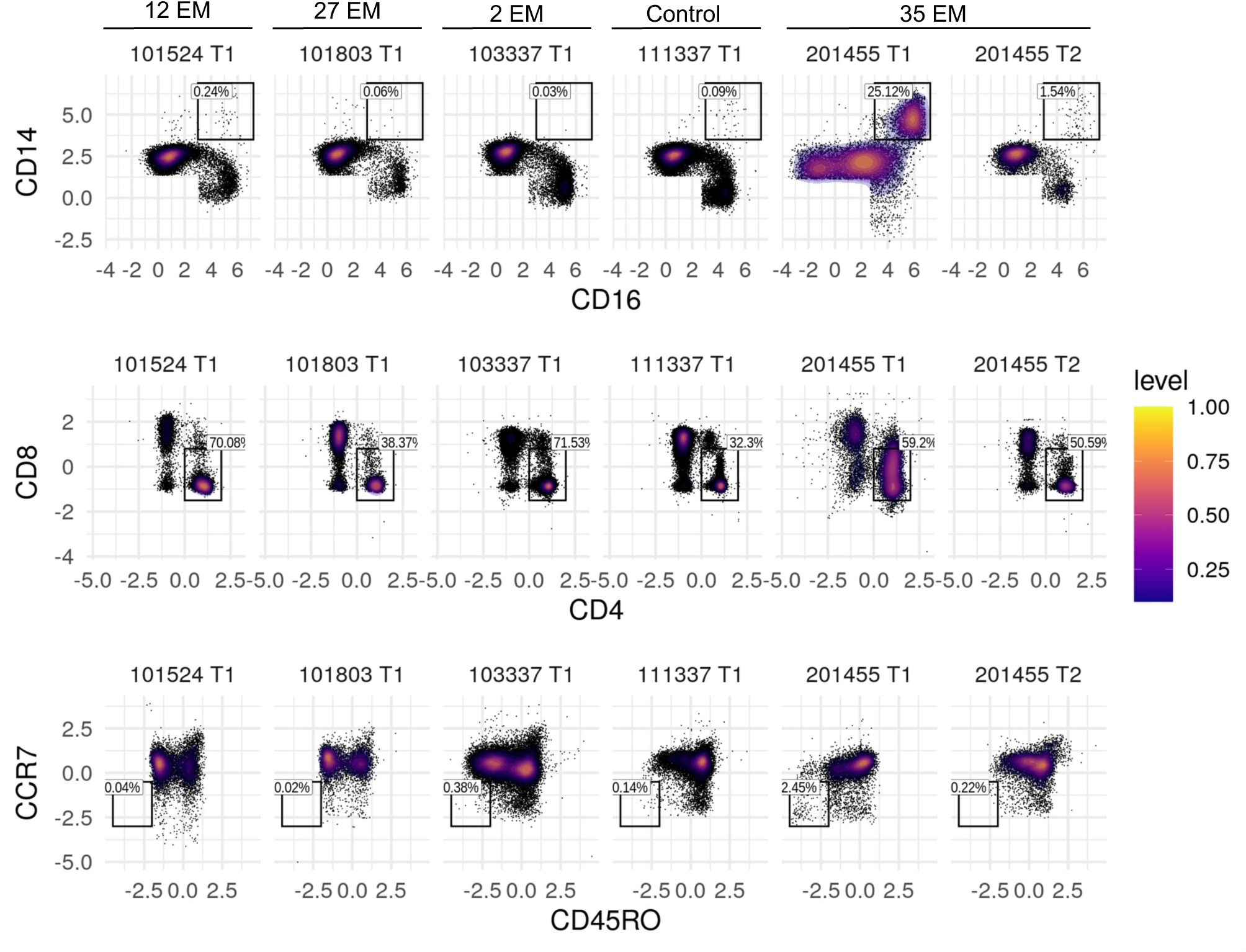
Pronounced PBMC changes only in a patient with 35 EM lesions. Representative flow cytometry plots of monocytes and T cells from patients with varying numbers of EM at diagnosis, and from a healthy control. Monocytes were defined as CD14^+^Dump^-^ and T cells were defined as CD3^+^ (parent gates not shown). Level indicates relative cell density.

**Figure S10:**
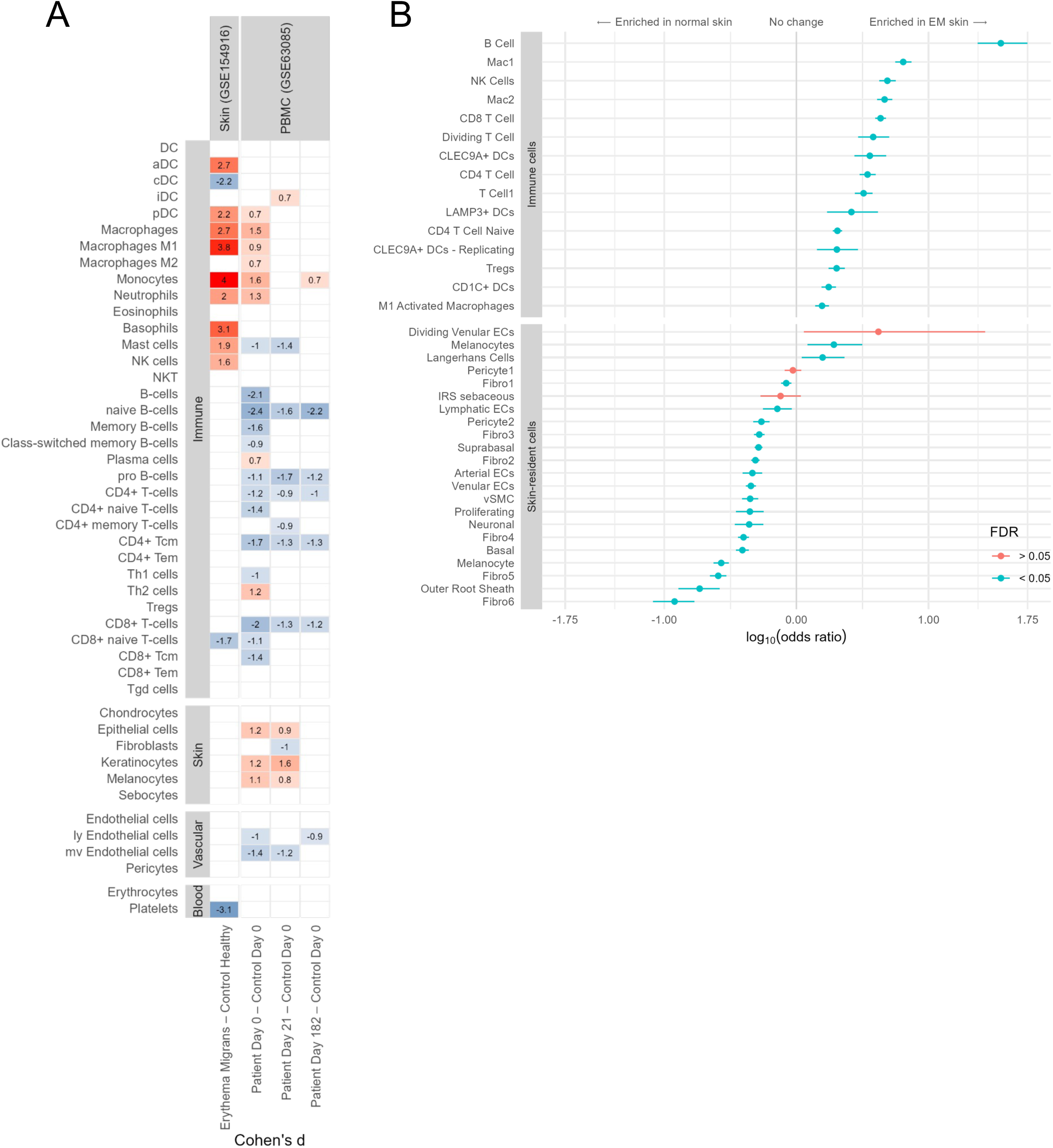
Immune and stromal cell abundance analyses in skin and PBMC transcriptomes from LD patients and controls. (A) Heatmap showing Cohen’s d effect sizes for cell type enrichment scores computed using xCell from publicly available bulk transcriptomic datasets of EM skin lesions (GSE154916) and PBMCs (GSE63085) from LD patients and controls. Positive values (red) indicate greater enrichment in patients; negative values (blue) indicate greater enrichment in controls. White boxes indicate statistically non-significant differences (p > 0.05). (B) Forest plot showing cell types with significantly different abundance in EM lesions versus unaffected skin, based on reanalysis of public skin scRNA-Seq data (GSE169440). Horizontal bars represent confidence intervals around odds ratios; a log10(odds ratio) of zero indicates no difference between groups. This panel resolves the broad cell type categories shown in Fig. 7b into finer subpopulations.

**Figure S11:**
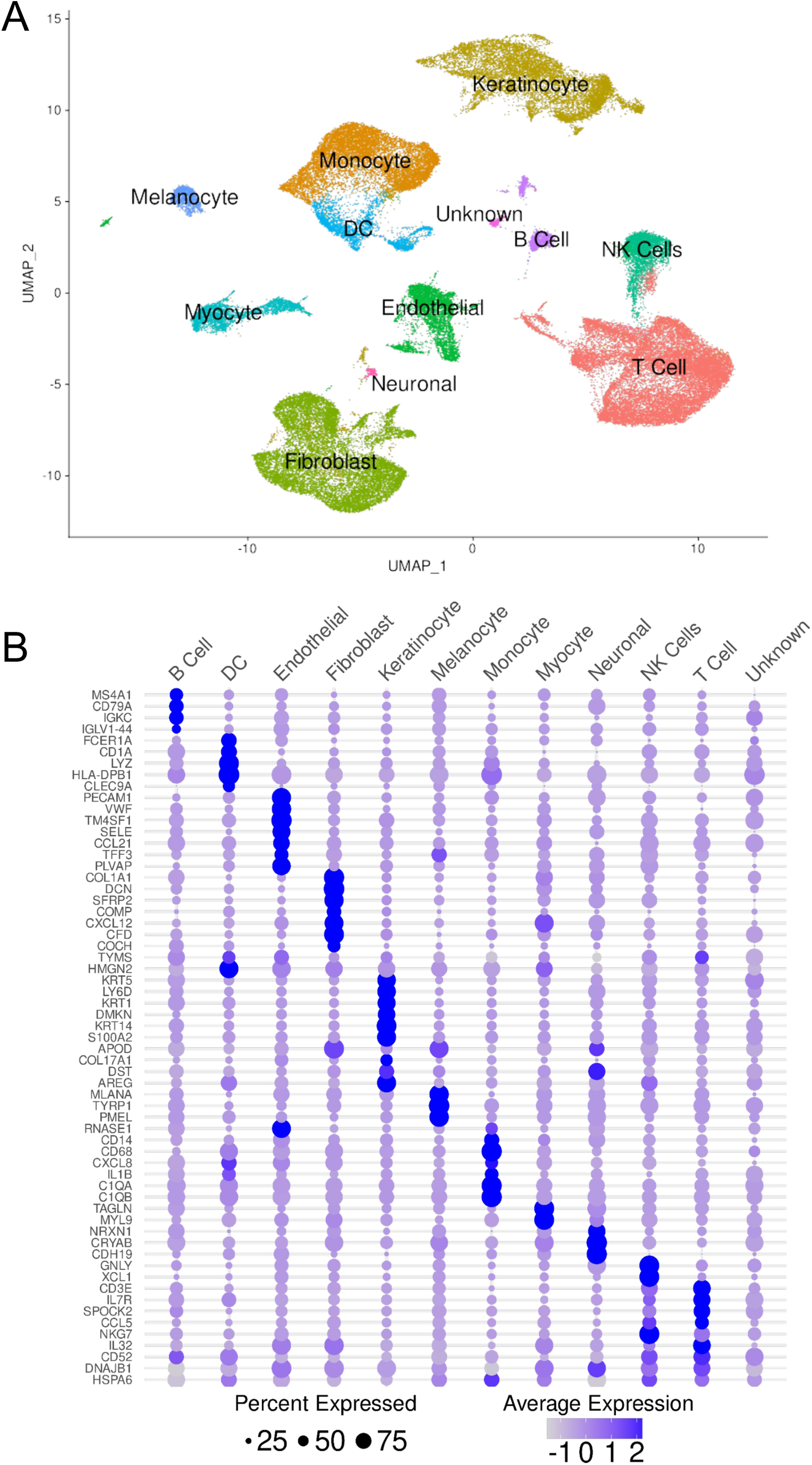
Analysis of scRNA-Seq data from EM skin lesions (GSE169440). **(A)** UMAP showing annotation of major cell populations at the site of infection. **(B)** Bubble plot showing marker genes (rows) for the cell clusters in A (columns). Dot size represents the average number of cells expressing each gene, and color represents the level of expression.

**Figure S12:**
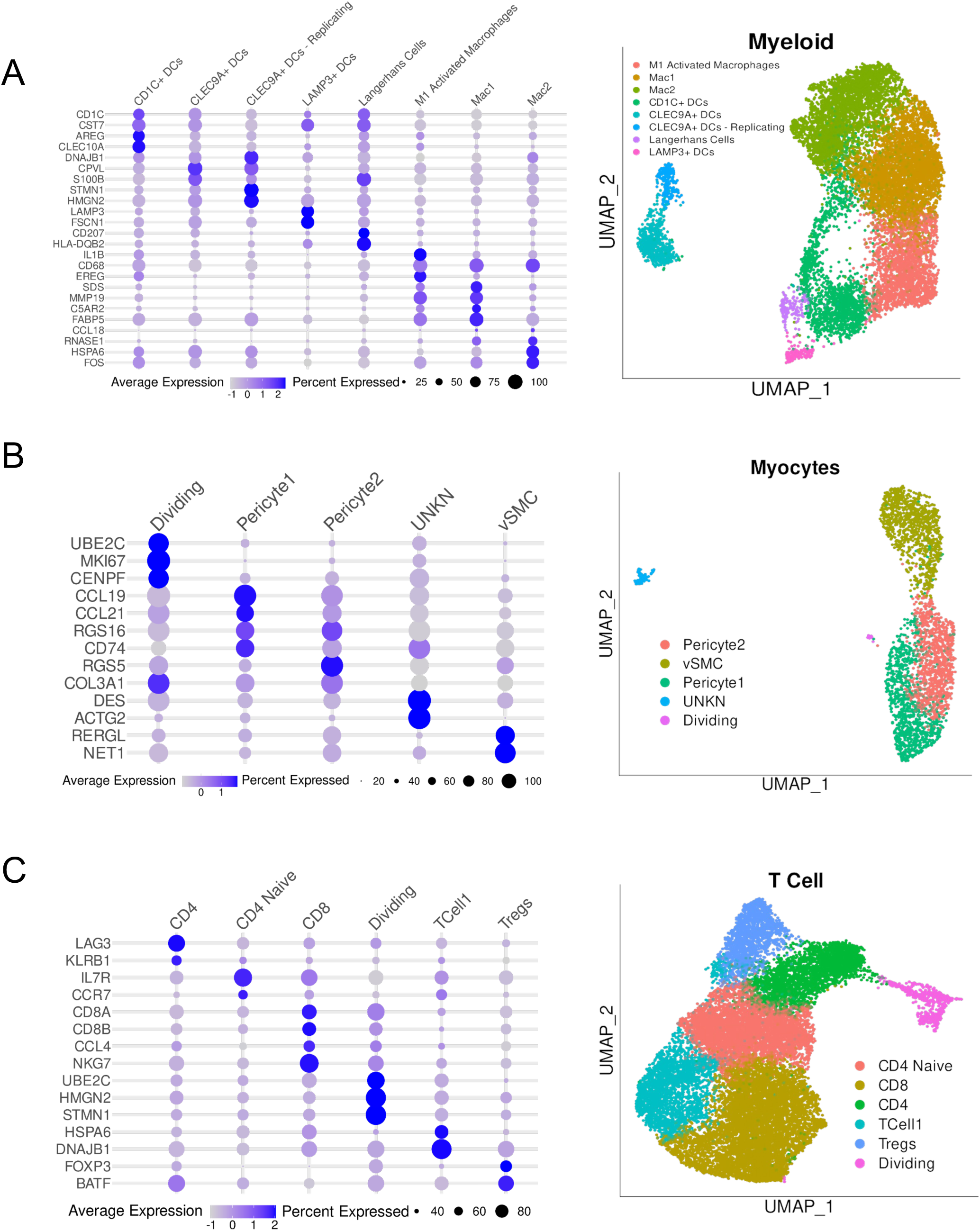
Sub-clustering of myeloid, myocyte, and T cell populations from Fig. S11. UMAP (right) and corresponding bubble plot (left) showing sub-clustering of myeloid **(A)**, myocyte **(B)**, and T cell **(C)** populations from Fig. S11. Dot size on bubble plots represents the average number of cells expressing each gene and color represents the level of expression.

**Figure S13:**
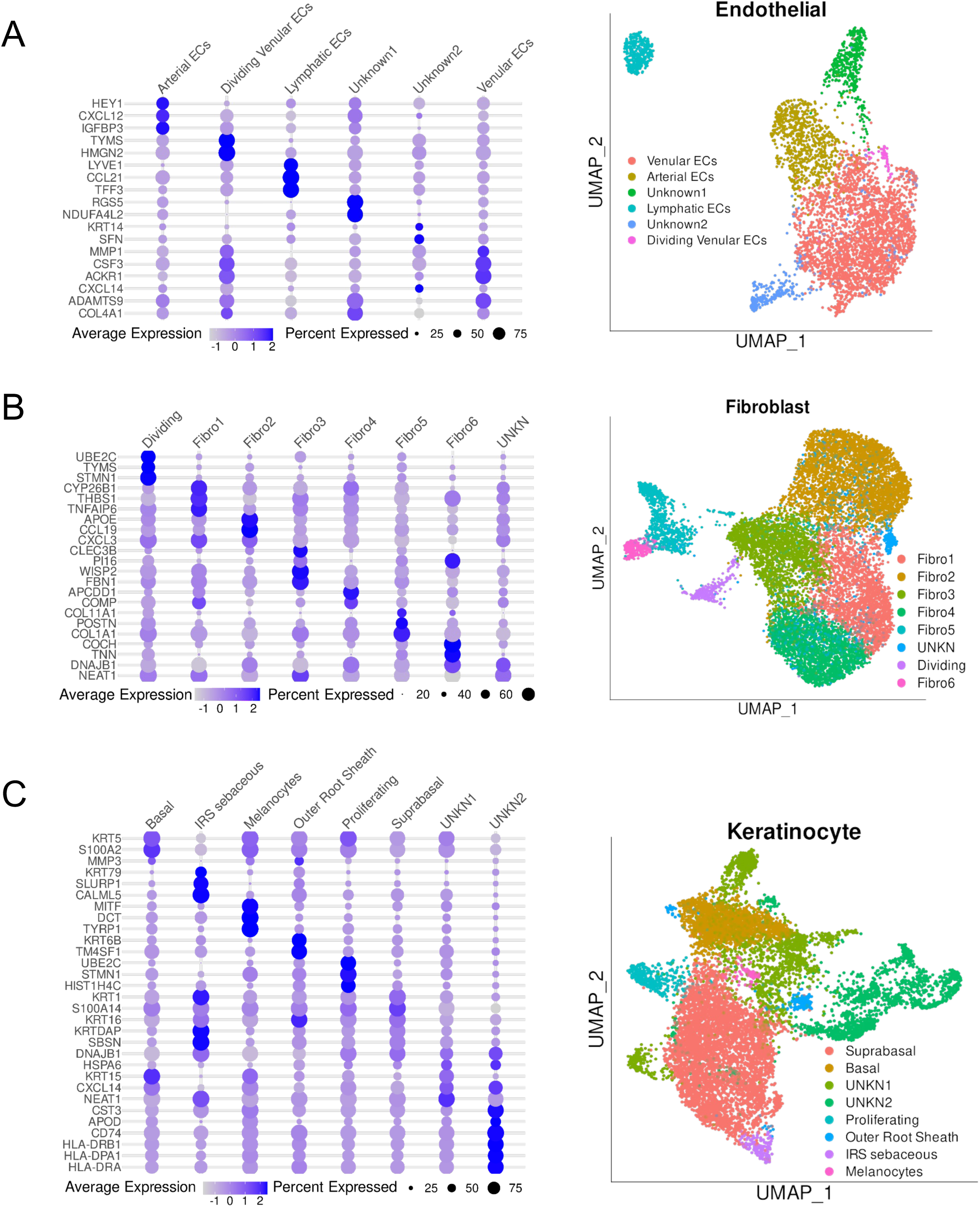
Sub-clustering of endothelial, fibroblast, and keratinocyte cell populations from Fig. S11. UMAP (right) and corresponding bubble plot (left) showing sub-clustering of endothelial **(A)**, fibroblast **(B)**, and keratinocyte **(C)** populations from Fig. S11. Dot size on bubble plots represents the average number of cells expressing each gene and color represents the level of expression.

## References

1. Mead, P.S. Epidemiology of Lyme disease. Infect Dis Clin North Am 29, 187–210 (2015).

2. Chaconas, G., Castellanos, M. & Verhey, T.B. Changing of the guard: How the Lyme disease spirochete subverts the host immune response. J Biol Chem 295, 301–313 (2020).

3. Voordouw, M.J., Lachish, S. & Dolan, M.C. The lyme disease pathogen has no effect on the survival of its rodent reservoir host. PLoS One 10, e0118265 (2015).

4. Yang, H., et al. By-degree Health and Economic Impacts of Lyme Disease, Eastern and Midwestern United States. Ecohealth 21, 56–70 (2024).

5. Gatewood, A.G., et al. Climate and tick seasonality are predictors of Borrelia burgdorferi genotype distribution. Appl Environ Microbiol 75, 2476–2483 (2009).

6. Shapiro, E.D. Clinical practice. Lyme disease. N Engl J Med 370, 1724–1731 (2014).

7. Steere, A.C. Lyme disease. N Engl J Med 345, 115–125 (2001).

8. Steere, A.C., et al. Lyme borreliosis. Nat Rev Dis Primers 2, 16090 (2016).

9. Stanek, G., Wormser, G.P., Gray, J. & Strle, F. Lyme borreliosis. Lancet 379, 461–473 (2012).

10. Wormser, G.P., et al. The clinical assessment, treatment, and prevention of lyme disease, human granulocytic anaplasmosis, and babesiosis: clinical practice guidelines by the Infectious Diseases Society of America. Clin Infect Dis 43, 1089–1134 (2006).

11. Rebman, A.W., et al. The Clinical, Symptom, and Quality-of-Life Characterization of a Well-Defined Group of Patients with Posttreatment Lyme Disease Syndrome. Front Med (Lausanne) 4, 224 (2017).

12. Choutka, J., Jansari, V., Hornig, M. & Iwasaki, A. Unexplained post-acute infection syndromes. Nat Med 28, 911–923 (2022).

13. Marques, A.R. Laboratory diagnosis of Lyme disease: advances and challenges. Infect Dis Clin North Am 29, 295–307 (2015).

14. Blum, L.K., et al. Robust B Cell Responses Predict Rapid Resolution of Lyme Disease. Front Immunol 9, 1634 (2018).

15. Kirpach, J., et al. Detection of a Low Level and Heterogeneous B Cell Immune Response in Peripheral Blood of Acute Borreliosis Patients With High Throughput Sequencing. Front Immunol 10, 1105 (2019).

16. Kelbauskas, L., Legutki, J.B. & Woodbury, N.W. Highly heterogenous humoral immune response in Lyme disease patients revealed by broad machine learning-assisted antibody binding profiling with random peptide arrays. Front Immunol 15, 1335446 (2024).

17. Sellati, T.J. & Barberio, D.M. Mechanisms of Dysregulated Antibody Response in Lyme Disease. Front Cell Infect Microbiol 10, 567252 (2020).

18. Hastey, C.J., Elsner, R.A., Barthold, S.W. & Baumgarth, N. Delays and diversions mark the development of B cell responses to Borrelia burgdorferi infection. J Immunol 188, 5612–5622 (2012).

19. Elsner, R.A., Hastey, C.J., Olsen, K.J. & Baumgarth, N. Suppression of Long-Lived Humoral Immunity Following Borrelia burgdorferi Infection. PLoS Pathog 11, e1004976 (2015).

20. Marques, A., et al. Transcriptome Assessment of Erythema Migrans Skin Lesions in Patients With Early Lyme Disease Reveals Predominant Interferon Signaling. J Infect Dis 217, 158–167 (2017).

21. Soloski, M.J., et al. Serum inflammatory mediators as markers of human Lyme disease activity. PLoS One 9, e93243 (2014).

22. Petzke, M.M., et al. Borrelia burgdorferi induces a type I interferon response during early stages of disseminated infection in mice. BMC Microbiol 16, 29 (2016).

23. Droyvold, W.B., Midthjell, K., Nilsen, T.I. & Holmen, J. Change in body mass index and its impact on blood pressure: a prospective population study. Int J Obes (Lond) 29, 650–655 (2005).

24. Clarke, D.J.B., et al. Predicting Lyme Disease From Patients’ Peripheral Blood Mononuclear Cells Profiled With RNA-Sequencing. Front Immunol 12, 636289 (2021).

25. Petzke, M.M., et al. Global Transcriptome Analysis Identifies a Diagnostic Signature for Early Disseminated Lyme Disease and Its Resolution. mBio 11(2020).

26. Knoops, B., Argyropoulou, V., Becker, S., Ferte, L. & Kuznetsova, O. Multiple Roles of Peroxiredoxins in Inflammation. Mol Cells 39, 60–64 (2016).

27. Xu, H., et al. New genetic and epigenetic insights into the chemokine system: the latest discoveries aiding progression toward precision medicine. Cell Mol Immunol 20, 739–776 (2023).

28. Andrea L. Wurster, M.J.G. Cytokines. in Encyclopedia of Biological Chemistry (ed. William J. Lennarz, M.D.L.) 550-555 (Elsevier, 2004).

29. Morris, R., Kershaw, N.J. & Babon, J.J. The molecular details of cytokine signaling via the JAK/STAT pathway. Protein Sci 27, 1984–2009 (2018).

30. Kanellis, J., et al. Stanniocalcin-1, an inhibitor of macrophage chemotaxis and chemokinesis. Am J Physiol Renal Physiol 286, F356–362 (2004).

31. Huang, L., et al. Anti-inflammatory and renal protective actions of stanniocalcin-1 in a model of anti-glomerular basement membrane glomerulonephritis. Am J Pathol 174, 1368–1378 (2009).

32. Yeung, B.H., Law, A.Y. & Wong, C.K. Evolution and roles of stanniocalcin. Mol Cell Endocrinol 349, 272–280 (2012).

33. Wang, J., et al. LSECtin interacts with BTN3A1 to inhibit T cell activation. The Journal of Immunology 204, 78.76–78.76 (2020).

34. Tang, L., et al. Liver sinusoidal endothelial cell lectin, LSECtin, negatively regulates hepatic T-cell immune response. Gastroenterology 137, 1498–1508 e1491-1495 (2009).

35. Jaakkola, P., et al. Targeting of HIF-alpha to the von Hippel-Lindau ubiquitylation complex by O2-regulated prolyl hydroxylation. Science 292, 468–472 (2001).

36. Celus, W., et al. Plexin-A4 Mediates Cytotoxic T-cell Trafficking and Exclusion in Cancer. Cancer Immunol Res 10, 126–141 (2022).

37. De Louche, C.D. & Roghanian, A. Human inhibitory leukocyte Ig-like receptors: from immunotolerance to immunotherapy. JCI Insight 7(2022).

38. Igarashi, K. & Watanabe-Matsui, M. Wearing red for signaling: the heme-bach axis in heme metabolism, oxidative stress response and iron immunology. Tohoku J Exp Med 232, 229–253 (2014).

39. Michels, A.A. & Bensaude, O. Hexim1, an RNA-controlled protein hub. Transcription 9, 262–271 (2018).

40. Asakura, A. Vascular endothelial growth factor gene regulation by HEXIM1 in heart. Circ Res 102, 398–400 (2008).

41. Kim, I., et al. Angiopoietin-1 regulates endothelial cell survival through the phosphatidylinositol 3’-Kinase/Akt signal transduction pathway. Circ Res 86, 24–29 (2000).

42. Potente, M., Gerhardt, H. & Carmeliet, P. Basic and therapeutic aspects of angiogenesis. Cell 146, 873–887 (2011).

43. David, L., et al. Bone morphogenetic protein-9 is a circulating vascular quiescence factor. Circ Res 102, 914–922 (2008).

44. Oksi, J., et al. Decreased interleukin-4 and increased gamma interferon production by peripheral blood mononuclear cells of patients with Lyme borreliosis. Infect Immun 64, 3620–3623 (1996).

45. Glickstein, L., et al. Inflammatory cytokine production predominates in early Lyme disease in patients with erythema migrans. Infect Immun 71, 6051–6053 (2003).

46. Mullegger, R.R., et al. Chemokine signatures in the skin disorders of Lyme borreliosis in Europe: predominance of CXCL9 and CXCL10 in erythema migrans and acrodermatitis and CXCL13 in lymphocytoma. Infect Immun 75, 4621–4628 (2007).

47. Fallahi, P., Elia, G. & Bonatti, A. Interferon-gamma-induced protein 10 in Lyme disease. Clin Ter 168, e146–e150 (2017).

48. Stanbery, A.G., Shuchi, S., Jakob von, M., Tait Wojno, E.D. & Ziegler, S.F. TSLP, IL-33, and IL-25: Not just for allergy and helminth infection. J Allergy Clin Immunol 150, 1302–1313 (2022).

49. Mohan, T., Deng, L. & Wang, B.Z. CCL28 chemokine: An anchoring point bridging innate and adaptive immunity. Int Immunopharmacol 51, 165–170 (2017).

50. Nies, J.F. & Panzer, U. IL-17C/IL-17RE: Emergence of a Unique Axis in T(H)17 Biology. Front Immunol 11, 341 (2020).

51. Strobl, J., et al. Tick feeding modulates the human skin immune landscape to facilitate tick-borne pathogen transmission. J Clin Invest 132(2022).

52. Kessler, O., et al. Semaphorin-3F is an inhibitor of tumor angiogenesis. Cancer Res 64, 1008–1015 (2004).

53. Pereira, M.G., et al. Angiotensin II-independent angiotensin-(1-7) formation in rat hippocampus: involvement of thimet oligopeptidase. Hypertension 62, 879–885 (2013).

54. Felbor, U., et al. Secreted cathepsin L generates endostatin from collagen XVIII. EMBO J 19, 1187–1194 (2000).

55. He, X., Zeng, H., Cantrell, A.C. & Chen, J.X. Regulatory role of TIGAR on endothelial metabolism and angiogenesis. J Cell Physiol 236, 7578–7590 (2021).

56. Krautter, F., et al. Galectin-9: A novel promoter of atherosclerosis progression. Atherosclerosis 363, 57–68 (2022).

57. Li, W., et al. Thymidine phosphorylase participates in platelet signaling and promotes thrombosis. Circ Res 115, 997–1006 (2014).

58. Dewberry, R., Holden, H., Crossman, D. & Francis, S. Interleukin-1 receptor antagonist expression in human endothelial cells and atherosclerosis. Arterioscler Thromb Vasc Biol 20, 2394–2400 (2000).

59. Fahey, E. & Doyle, S.L. IL-1 Family Cytokine Regulation of Vascular Permeability and Angiogenesis. Front Immunol 10, 1426 (2019).

60. Mitchell, A., et al. LILRA5 is expressed by synovial tissue macrophages in rheumatoid arthritis, selectively induces pro-inflammatory cytokines and IL-10 and is regulated by TNF-alpha, IL-10 and IFN-gamma. Eur J Immunol 38, 3459-3473 (2008).

61. Fan, X., et al. Paired immunoglobulin-like receptor B regulates platelet activation. Blood 124, 2421–2430 (2014).

62. Xue, C., et al. Tryptophan metabolism in health and disease. Cell Metab 35, 1304–1326 (2023).

63. Halperin, J.J. & Heyes, M.P. Neuroactive kynurenines in Lyme borreliosis. Neurology 42, 43–50 (1992).

64. Wickstrom, R., et al. The Kynurenine Pathway is Differentially Activated in Children with Lyme Disease and Tick-Borne Encephalitis. Microorganisms 9(2021).

65. Elfenbein, A. & Simons, M. Syndecan-4 signaling at a glance. J Cell Sci 126, 3799–3804 (2013).

66. Baeyens, N., et al. Syndecan 4 is required for endothelial alignment in flow and atheroprotective signaling. Proc Natl Acad Sci U S A 111, 17308–17313 (2014).

67. Xiong, Y. & Hla, T. S1P control of endothelial integrity. Curr Top Microbiol Immunol 378, 85–105 (2014).

68. Zhang, Y., et al. CCL17 acts as a novel therapeutic target in pathological cardiac hypertrophy and heart failure. J Exp Med 219(2022).

69. Wu, B., et al. Crk and Crkl Are Required in the Endocardial Lineage for Heart Valve Development. J Am Heart Assoc 12, e029683 (2023).

70. Choi, S.H., et al. Dickkopf-1 induces angiogenesis via VEGF receptor 2 regulation independent of the Wnt signaling pathway. Oncotarget 8, 58974–58984 (2017).

71. Park, M.H., Sung, E.A., Sell, M. & Chae, W.J. Dickkopf1: An Immunomodulator in Tissue Injury, Inflammation, and Repair. Immunohorizons 5, 898–908 (2021).

72. Barreto, J., Karathanasis, S.K., Remaley, A. & Sposito, A.C. Role of LOX-1 (Lectin-Like Oxidized Low-Density Lipoprotein Receptor 1) as a Cardiovascular Risk Predictor: Mechanistic Insight and Potential Clinical Use. Arterioscler Thromb Vasc Biol 41, 153–166 (2021).

73. Sayers, E.W., et al. GenBank. Nucleic Acids Res 50, D161–D164 (2022).

74. Kim, Y.H., et al. A MST1-FOXO1 cascade establishes endothelial tip cell polarity and facilitates sprouting angiogenesis. Nat Commun 10, 838 (2019).

75. Marfia, G., et al. Decreased serum level of sphingosine-1-phosphate: a novel predictor of clinical severity in COVID-19. EMBO Mol Med 13, e13424 (2021).

76. Houten, S.M., Violante, S., Ventura, F.V. & Wanders, R.J. The Biochemistry and Physiology of Mitochondrial Fatty Acid beta-Oxidation and Its Genetic Disorders. Annu Rev Physiol 78, 23–44 (2016).

77. Fitzgerald, B.L., et al. Host Metabolic Response in Early Lyme Disease. J Proteome Res 19, 610–623 (2020).

78. Molins, C.R., et al. Metabolic differentiation of early Lyme disease from southern tick-associated rash illness (STARI). Sci Transl Med 9(2017).

79. Weis, E.M., et al. Ketone body oxidation increases cardiac endothelial cell proliferation. EMBO Mol Med 14, e14753 (2022).

80. Patella, F., et al. Proteomics-based metabolic modeling reveals that fatty acid oxidation (FAO) controls endothelial cell (EC) permeability. Mol Cell Proteomics 14, 621–634 (2015).

81. Cruz, M.M., et al. Palmitoleic acid (16:1n7) increases oxygen consumption, fatty acid oxidation and ATP content in white adipocytes. Lipids Health Dis 17, 55 (2018).

82. de Souza, C.O., et al. Palmitoleic Acid has Stronger Anti-Inflammatory Potential in Human Endothelial Cells Compared to Oleic and Palmitic Acids. Mol Nutr Food Res 62, e1800322 (2018).

83. Schillemans, M., Karampini, E., Kat, M. & Bierings, R. Exocytosis of Weibel-Palade bodies: how to unpack a vascular emergency kit. J Thromb Haemost 17, 6–18 (2019).

84. Silva, E.E., Skon-Hegg, C., Badovinac, V.P. & Griffith, T.S. The Calm after the Storm: Implications of Sepsis Immunoparalysis on Host Immunity. J Immunol 211, 711–719 (2023).

85. Aran, D., Hu, Z. & Butte, A.J. xCell: digitally portraying the tissue cellular heterogeneity landscape. Genome Biol 18, 220 (2017).

86. Jiang, R., et al. Single-cell immunophenotyping of the skin lesion erythema migrans identifies IgM memory B cells. JCI Insight 6 (2021).

87. Sugita, S., et al. T-cell suppression by programmed cell death 1 ligand 1 on retinal pigment epithelium during inflammatory conditions. Invest Ophthalmol Vis Sci 50, 2862–2870 (2009).

88. Le, C.T., et al. Regulation of human and mouse bystander T cell activation responses by PD-1. JCI Insight 8(2023).

89. Bockenstedt, L.K., et al. What ticks do under your skin: two-photon intravital imaging of Ixodes scapularis feeding in the presence of the lyme disease spirochete. Yale J Biol Med 87, 3–13 (2014).

90. Jabbari, N., et al. Whole genome sequence and comparative analysis of Borrelia burgdorferi MM1. PLoS One 13, e0198135 (2018).

91. Kapellos, T.S., et al. Human Monocyte Subsets and Phenotypes in Major Chronic Inflammatory Diseases. Front Immunol 10, 2035 (2019).

92. Libby, P. Endothelial inflammation in COVID-19. Science 386, 972–973 (2024).

93. Ng, C.T., Fong, L.Y. & Abdullah, M.N.H. Interferon-gamma (IFN-gamma): Reviewing its mechanisms and signaling pathways on the regulation of endothelial barrier function. Cytokine 166, 156208 (2023).

94. Asanuma, K., et al. Soluble programmed death-ligand 1 rather than PD-L1 on tumor cells effectively predicts metastasis and prognosis in soft tissue sarcomas. Sci Rep 10, 9077 (2020).

95. Suciu-Foca, N., et al. Soluble Ig-like transcript 3 inhibits tumor allograft rejection in humanized SCID mice and T cell responses in cancer patients. J Immunol 178, 7432–7441 (2007).

96. Love, A.C., Schwartz, I. & Petzke, M.M. Induction of indoleamine 2,3-dioxygenase by Borrelia burgdorferi in human immune cells correlates with pathogenic potential. J Leukoc Biol 97, 379–390 (2015).

97. Baban, B., et al. IDO activates regulatory T cells and blocks their conversion into Th17-like T cells. J Immunol 183, 2475–2483 (2009).

98. Stone, T.W. & Williams, R.O. Modulation of T cells by tryptophan metabolites in the kynurenine pathway. Trends Pharmacol Sci 44, 442–456 (2023).

99. Mezrich, J.D., et al. An interaction between kynurenine and the aryl hydrocarbon receptor can generate regulatory T cells. J Immunol 185, 3190–3198 (2010).

100. Longhi, S.A., et al. Cytokine production but lack of proliferation in peripheral blood mononuclear cells from chronic Chagas’ disease cardiomyopathy patients in response to T. cruzi ribosomal P proteins. PLoS Negl Trop Dis 8, e2906 (2014).

101. Johnson, W.E., Li, C. & Rabinovic, A. Adjusting batch effects in microarray expression data using empirical Bayes methods. Biostatistics 8, 118–127 (2007).

102. Ritchie, M.E., et al. limma powers differential expression analyses for RNA-sequencing and microarray studies. Nucleic Acids Res 43, e47 (2015).

103. Benjamini, Y. & Hochberg, Y. Controlling the False Discovery Rate: A Practical and Powerful Approach to Multiple Testing. Journal of the Royal Statistical Society: Series B (Methodological) 57, 289–300 (1995).

104. Wu, T., et al. clusterProfiler 4.0: A universal enrichment tool for interpreting omics data. Innovation (Camb) 2, 100141 (2021).

105. Liberzon, A., et al. Molecular signatures database (MSigDB) 3.0. Bioinformatics 27, 1739–1740 (2011).

106. Friedman, J., Hastie, T. & Tibshirani, R. Regularization Paths for Generalized Linear Models via Coordinate Descent. J Stat Softw 33, 1–22 (2010).

107. Groll, A.T., G. Variable Selection for Generalized Linear Mixed Models by L1-Penalized Estimation. Statistics and Computing 24, 137–154 (2014).

108. Evans, A.M., DeHaven, C.D., Barrett, T., Mitchell, M. & Milgram, E. Integrated, nontargeted ultrahigh performance liquid chromatography/electrospray ionization tandem mass spectrometry platform for the identification and relative quantification of the small-molecule complement of biological systems. Anal Chem 81, 6656–6667 (2009).

109. Zeileis, A.H., T. Diagnostic Checking in Regression Relationships. R News 2, 7–10 (2002).

110. Picart-Armada, S., Fernandez-Albert, F., Vinaixa, M., Yanes, O. & Perera-Lluna, A. FELLA: an R package to enrich metabolomics data. BMC Bioinformatics 19, 538 (2018).

111. Hahne, F., et al. flowCore: a Bioconductor package for high throughput flow cytometry. BMC Bioinformatics 10, 106 (2009).

112. Malek, M., et al. flowDensity: reproducing manual gating of flow cytometry data by automated density-based cell population identification. Bioinformatics 31, 606–607 (2015).

113. Mouselimis, L.e.a. ClusterR: Gaussian Mixture Models, K-Means, Mini-Batch-Kmeans, K-Medoids and Affinity Propagation Clustering. (2022).

114. Azad, A., Rajwa, B. & Pothen, A. flowVS: channel-specific variance stabilization in flow cytometry. BMC Bioinformatics 17, 291 (2016).

115. Fletez-Brant, K., Spidlen, J., Brinkman, R.R., Roederer, M. & Chattopadhyay, P.K. flowClean: Automated identification and removal of fluorescence anomalies in flow cytometry data. Cytometry A 89, 461–471 (2016).

116. Lux, M., et al. flowLearn: fast and precise identification and quality checking of cell populations in flow cytometry. Bioinformatics 34, 2245–2253 (2018).

117. Nowicka, M., et al. CyTOF workflow: differential discovery in high-throughput high-dimensional cytometry datasets. F1000Res 6, 748 (2017).

118. Van Gassen, S., et al. FlowSOM: Using self-organizing maps for visualization and interpretation of cytometry data. Cytometry A 87, 636–645 (2015).

119. Wilkerson, M.D. & Hayes, D.N. ConsensusClusterPlus: a class discovery tool with confidence assessments and item tracking. Bioinformatics 26, 1572–1573 (2010).

120. Weber, L.M., Nowicka, M., Soneson, C. & Robinson, M.D. diffcyt: Differential discovery in high-dimensional cytometry via high-resolution clustering. Commun Biol 2, 183 (2019).

121. Buuren, S.v.G.-O., K. mice: Multivariate Imputation by Chained Equations in R. J. Stat. Softw. 45, 1–67 (2011).

122. Huang, Y., McCarthy, D.J. & Stegle, O. Vireo: Bayesian demultiplexing of pooled single-cell RNA-seq data without genotype reference. Genome Biol 20, 273 (2019).

123. McCarthy, D.J., Campbell, K.R., Lun, A.T. & Wills, Q.F. Scater: pre-processing, quality control, normalization and visualization of single-cell RNA-seq data in R. Bioinformatics 33, 1179–1186 (2017).

124. Stuart, T., et al. Comprehensive Integration of Single-Cell Data. Cell 177, 1888–1902 e1821 (2019).

125. Hao, Y., et al. Integrated analysis of multimodal single-cell data. Cell 184, 3573–3587 e3529 (2021).

126. Xu, C. & Su, Z. Identification of cell types from single-cell transcriptomes using a novel clustering method. Bioinformatics 31, 1974–1980 (2015).

127. Blondel, V.D.J.-L.G.R.L.a.E.L. Fast Unfolding of Communities in Large Networks. Journal of Statistical Mechanics: Theory and Experiment 10(2008).

128. Aran, D., et al. Reference-based analysis of lung single-cell sequencing reveals a transitional profibrotic macrophage. Nat Immunol 20, 163–172 (2019).

129. Monaco, G., et al. RNA-Seq Signatures Normalized by mRNA Abundance Allow Absolute Deconvolution of Human Immune Cell Types. Cell Rep 26, 1627–1640 e1627 (2019).

130. Squair, J.W., et al. Confronting false discoveries in single-cell differential expression. Nat Commun 12, 5692 (2021).

131. Love, M.I., Huber, W. & Anders, S. Moderated estimation of fold change and dispersion for RNA-seq data with DESeq2. Genome Biol 15, 550 (2014).

132. Aucott, J.N., et al. CCL19 as a Chemokine Risk Factor for Posttreatment Lyme Disease Syndrome: a Prospective Clinical Cohort Study. Clin Vaccine Immunol 23, 757–766 (2016).

133. Davis, S. & Meltzer, P.S. GEOquery: a bridge between the Gene Expression Omnibus (GEO) and BioConductor. Bioinformatics 23, 1846–1847 (2007).

134. Mahi, N.A., Najafabadi, M.F., Pilarczyk, M., Kouril, M. & Medvedovic, M. GREIN: An Interactive Web Platform for Re-analyzing GEO RNA-seq Data. Sci Rep 9, 7580 (2019).

135. Law, C.W., Chen, Y., Shi, W. & Smyth, G.K. voom: Precision weights unlock linear model analysis tools for RNA-seq read counts. Genome Biol 15, R29 (2014).

136. Price, N.D., et al. A wellness study of 108 individuals using personal, dense, dynamic data clouds. Nat Biotechnol 35, 747–756 (2017).

137. Csardi, G., & Nepusz, T. The Igraph Software Package for Complex Network Research. *InterJournal*, Complex Systems 1695, 1–9 (2006).

